# Gasdermin B over-expression arbitrates HER2-targeted therapy resistance by inducing protective autophagy

**DOI:** 10.1101/2021.07.01.450506

**Authors:** Manuel Gámez-Chiachio, Ángela Molina-Crespo, Carmen Ramos-Nebot, Jeannette Martinez-Val, Lidia Martinez, Katja Gassner, Francisco J. Llobet, Claudia Gonzalo-Consuegra, Marco Cordani, Cristina Bernadó-Morales, Eva Diaz, Alejandro Rojo-Sebastian, Juan Carlos Triviño, Laura Sanchez, Ruth Rodríguez-Barrueco, Joaquín Arribas, David Llobet-Navás, David Sarrió, Gema Moreno-Bueno

**Affiliations:** Departamento de Bioquímica, Universidad Autónoma de Madrid (UAM), Instituto de Investigaciones Biomédicas ‘Alberto Sols’ (CSIC-UAM), IdiPAZ, Madrid, Spain; Centro de Investigación Biomédica en Red de Cáncer (CIBERONC), Instituto de Salud Carlos III, Madrid, Spain; Departamento de Zoología, Genética, Antropología Física, Universidad Santiago de Compostela, Lugo, Spain; Mecanismos Moleculares y Terapía Experimental en Oncologia-Programa Oncobell, Idibell – L’Hospitalet de Llobregat, Spain; Programa de Investigación Preclínica, Vall d’Hebron Institute of Oncology (VHIO), Barcelona, Spain; Fundación MD Anderson Internacional, Madrid, Spain; MD Anderson Cancer Center, Madrid, Spain; Sistemas Genómicos, Paterna, Valencia, Spain; Unidad de Anatomia, Departamento de Patologia y Terapèutica Experimental, Facultad de Medicina, Universidad de Barcelona (UB) – L’Hospitalet de Llobregat, Spain; Programa de Investigación en Cáncer, IMIM (Hospital del Mar Medical Research Institute), Barcelona, Spain; Institució Catalana de Recerca i Estudis Avançats (ICREA), Barcelona, Spain

**Keywords:** Gasdermin B, protective autophagy, anti-HER2 therapy, drug resistance, HER2 breast cancer, gastroesophageal tumors, LC3B

## Abstract

**Purpose:** Gasdermin B (GSDMB) over-expression promotes poor prognosis and aggressive behavior in HER2 breast cancer by increasing cell invasion, metastasis and resistance to therapy. Decoding the molecular mechanism of GSDMB-mediated drug resistance is crucial to identify novel effective targeted treatments for HER2/GSDMB aggressive tumors.

**Experiment design:** To decipher the functional relevance of GSDMB in promoting resistance to HER2-targeted therapies we performed several molecular approaches (immunoblot, qRT-PCR, flow cytometry, immunoprecipitation and confocal microscopy) in different breast and gastric carcinoma cell models. The results were confirmed in Patient Derived Xenografts (PDX) by qRT-PCR and in clinical human cancer samples by immunohistochemistry. Finally, we validated the efficacy of the identified targeted treatment in HER2/GSDMB cancers using two complementary *in vivo* preclinical models (tumor xenografts in mice and zebrafish).

**Results:** We discovered that GSDMB up-regulation renders HER2 breast and gastric cancer cells more resistant to anti-HER2 agents by promoting protective autophagy. Consistent with this, we proved that the combination of lapatinib with the autophagy inhibitor chloroquine increases the therapeutic response specifically in GSDMB-positive tumors *in vitro* and *in vivo* using zebrafish and mice preclinical cancer models. Mechanistically, we confirmed that the GSDMB N-terminal domain interacts with the autophagy protein LC3B. Finally, we validated these results in clinical samples of breast and gastric cancers, where GSDMB/LC3B co-expression associates significantly with relapse.

**Conclusion:** Our findings uncovered a novel functional link between GSDMB over-expression and LC3B-mediated protective autophagy in response to HER2-targeted therapies and provide a new and accessible therapeutic approach for HER2/GSDMB+ cancers with adverse clinical outcome.

**TRANSLATIONAL RELEVANCE:** Identifying the biomarkers and mechanisms of therapy resistance is a main challenge in current oncology. In this regard, Gasdermin-B (GSDMB) over-expression, which was initially found in >60% HER2 breast cancers, promotes resistance to therapy through an unknown molecular mechanism. In the present work, we revealed for the first time that in HER2 gastric and breast cancers GSDMB mediates innate and acquired resistance to HER2-targeted drugs through the promotion of a pro-survival autophagy mechanism that requires the interaction of GSDMB with LC3B. Accordingly, GSDMB/LC3B co-expression in human breast and gastric cancer clinical samples associates with relapse. To reverse this anti-drug effect, we developed a therapeutic approach based on the combination of the autophagy inhibitor chloroquine with lapatinib that showed significant efficacy both *in vitro* and *in vivo* on GSDMB-positive tumors. Our findings provide an accessible (FDA-approved drugs) therapeutic combination to treat effectively HER2/GSDMB over-expressing tumors with poor clinical outcome.

## INTRODUCTION

*HER2* amplification/overexpression occurs frequently in breast (1) and gastric cancers (2). While the association between HER2 status and poor prognosis in breast cancer is well-established, the prognostic impact of HER2 positivity in gastric tumors is still open to debate (3). Although different targeted anti-HER2 therapies have been developed (4), including HER2 blocking antibodies (e.g, trastuzumab, pertuzumab, T-DM1), and tyrosine kinase inhibitors (such as lapatinib), many patients develop drug resistance (4). A variety of distinct tumor resistance mechanisms to these agents have been proposed (5), that individually can explain a limited set of cases. In this regard, our group has proved that Gasdermin-B (GSDMB) over-expression and/or amplification, which occurs in more than 60% of HER2 breast cancers, is significantly associated with poor prognosis and reduced trastuzumab response independently of the hormone receptor status or histological grade (6). Moreover, GSDMB overexpression promotes cell motility, invasion and metastasis in breast cancer cell lines (7,8) but it does not affect cell proliferation (7). Interestingly, our recent data have shown that GSDMB over-expression is a novel therapeutic target in HER2 breast tumors (8), since the intracellular delivery of an GSDMB antibody using nanoparticles significantly reduces the tumor growth and metastasis development in HER2 breast tumors by inducing cancer cell death *in vivo* (8).

GSDMB is one of the six Gasdermin (GSDM) genes in the human genome (along with GSDMA, C, D, GSDME/DFNA5 and DFNB59/PJVK) (9). It is also frequently over-expressed in liver, colon, cervical cancers and in gastric carcinomas where GSDMB upregulation associates with tumor invasion (reviewed in 10). In addition to mediating diverse pro-tumor or anti-tumor effects (reviewed in (10)), recent data suggest that all GSDMs, under specific circumstances, can trigger a lytic and pro-inflammatory cell death mechanism, known as pyroptosis (11,12). This pro-cell death function is normally auto-inhibited through the intramolecular interaction of GSDM N-terminal and C-terminal domains, but upon cleavage by specific caspases and other proteases, the released N-terminal domain forms membrane pores that subsequently lead to cell lysis (reviewed in 19,20). In the case of GSDMB, the pro-cell death function under physiological conditions is a matter of intense debate (15–18) although a recent study has revealed that lymphocyte-derived granzyme A can cleave GSDMB within tumor cells, thus provoking a pyroptotic cancer cell death (19). In this sense, an activated antitumor immune response mediated by NK and T-cytotoxic cells could reduce tumor growth of murine colon and melanoma cells exogenously overexpressing GSDMB (19).

Given the aforementioned evidences demonstrating a role for GSDMB in tumor biology and clinical response, here we have unraveled the functional mechanism by which GSDMB over-expression can promote resistance to anti-HER2 therapies in human HER2 breast and gastric breast cancer cells. We show that GSDMB increases pro-survival autophagy likely through its cooperation with the microtubule-associated protein light chain 3B (LC3B), an important modulator of the autophagic machinery (20,21). Protective autophagy has emerged as a new resistance mechanism to diverse HER2-targeted drugs (22,23). In fact, the autophagy inhibition has been proposed as a new mechanism to resistance abrogation in HER2+ tumors (24) although the molecular mechanisms involved are currently unrevealed. Interestingly, we demonstrated that the combination of anti-HER2 treatment with the autophagy inhibitor chloroquine (CQ) increases the therapeutic response specifically in GSDMB-positive tumors *in vitro* and *in vivo* using different preclinical models (zebrafish and mice). Overall, we have uncovered a novel functional link between GSDMB over-expression and LC3B-mediated protective autophagy response to HER2-targeted therapies. This work provides a new therapeutic approach for HER2/GSDMB+ cancers with poor clinical outcome.

## MATERIAL AND METHODS

### Human tumor samples

The present study was carried out using a HER2 breast carcinoma series which were previously reported (6) and a HER2 intestinal gastric tumor cohort. Gastric tumors were acquired from the Biobank of the Anatomy Pathology Department (record number B.0000745, ISCIII National Biobank network) of the MD Anderson Cancer Center Madrid, Madrid, Spain. The mean patient age at diagnosis was 59.8±12.7 years (range, 29 to 85 years), 24.2% of tumors were diagnosed in women. All tumors were diagnostic as grade 2 or 3. According to the TNM classification staging, 11 were stage II, and 22 were stage III-IV. HER2 staining was performed at the diagnosis according to the established protocols (25). Immunohistological and clinical data of these tumors are provided in **Supplementary Table 1**. This study was performed following standard ethical procedures of the Spanish regulation (Ley de Investigación Orgánica Biomédica, 14 July 2007) and was approved by the ethic committees of the MD Anderson Cancer Center Madrid, Madrid, Spain.

### Cell culture

HCC1954, NCI-N87 and HEK293T cell lines were obtained from the American Type Cell Culture (ATCC) and OE19 cell line from the Deutsche Sammlung von Mikroorganismen und Zellkulturen (DSMZ). Cells were cultured following the supplier conditions. Cells were authenticated by STR-profiling according to ATCC or DSMZ guidelines. In order to generate lapatinib resistant (named LR) cell lines, both HCC1954 and OE19 parental cells were cultured up to 6 months in the presence of increasing concentrations of lapatinib from 0.2 μM up to 2 μM (HCC1954 cells, corresponds to IC50) or 1.5 μM (OE19, IC70). Resistant status was analyzed by cell viability assays using AlamarBlue (Bio-Rad), according to the manufacturer’s protocol. Then, HCC1954 LR and OE19 LR cells were cultured in the continuous presence of 2 μM and 1.5 μM lapatinib, respectively. In parallel, HCC1954 and OE19 control cells (C) were generated by chronic treatment with the vehicle DMSO (same amount than the corresponding lapatinib-resistant cells). The cell culture establishment from HER2-positive Patient Derived Xenografts (PDXs) was performed following the protocol previously described (26).

### Animal *in vivo* studies

All the experimental procedures with mice were approved by the internal ethical research and animal welfare committee (IDIBELL and IIB, UAM), and by the Local Authorities (Generalitat Catalana, B-9900010 and Comunidad de Madrid, PROEX424/15, respectively). They complied with the European Union (Directive 2010/63/UE) and Spanish Government guidelines (Real Decreto 53/20133). Furthermore, zebrafish studies (AB strain, *Danio rerio*) were performed with the agreement of the Bioethics Committee for animal experimentation of the University of Santiago of Compostela (CEEA-LU), REGA code: ES270280346401. The detailed description of the different *in vivo* experiments is provided in Supplementary Methods.

### Statistics and reproducibility

The Chi-square contingency test with Yates correction, or Fisher’s exact test, was used to determine the statistical significance of the relationships between immunohistochemical and clinico-pathological features. GraphPad Prism software was used for graphic representation and statistical analysis. Error bars represent the mean ± s.e.m of at least three independent experiments. Data were tested for normality, and paired sets of data were compared using paired Student’s t-test (two-tailed).

## RESULTS

### Anti-HER2 therapies up-regulate GSDMB expression and high levels of GSDMB protein associates with resistance to these drugs

GSDMB over-expression is a maker of poor prognosis associated with trastuzumab resistance in HER2 breast carcinoma patients in both neoadjuvant and adjuvant treatment settings (6). Moreover, high levels of GSDMB decrease sensitivity to trastuzumab *in vitro* in HCC1954 and SKBR3 cells, and its expression increases during the acquisition of trastuzumab resistance in HER2+ breast cancer PDX models (6). Here, to address whether GSDMB plays also a role in the clinical behavior of HER2 gastric tumors, we first analyzed GSDMB expression in a cohort of 33 tumors, and observed strong cytoplasmic and nuclear GSDMB staining in almost 60% of HER2 gastric tumors (Supplementary Fig. S1, see Supplementary Table S1 for clinical and molecular description of the tumor series). Similar to HER2 breast tumors (6), GSDMB over-expression in gastric carcinoma is significantly associated with the presence of distant metastasis (14/18, p= 0.032) or show a strong tendency with relapse (13/18, p=0.060, Supplementary Fig. S1B), thus supporting the relationship between high levels of GSDMB and poor prognosis in gastric tumors. Next, to test if in breast and gastroesophageal cancers GSDMB is functionally involved in regulating drug response/resistance also to the HER2 tyrosine kinase inhibitor lapatinib, we used three HER2+ cancer cell lines that endogenously express GSDMB, HCC1954 (breast cancer), OE19 (esophageal) and NCI-N87 (gastric). First, we treated these cells for different time points (up to 72 h) with their corresponding IC50 of lapatinib (**Fig. 1A** and **B**). Additionally, for comparison, trastuzumab treatment was carried out only in OE19 and NCI-N87 cells (Supplementary Fig. S2A and S2B), since HCC1954 cells are intrinsically highly resistant to this drug (8,27). Both, lapatinib (**Fig. 1A** and **B**) and trastuzumab (Supplementary Fig. S2A and S2B) treatments provoke a sharp induction of GSDMB mRNA, which peaked at 24-48 h, in all tested models. HER2 upregulation was also detected as previously reported (28). At the protein level, while GSDMB was strongly upregulated by lapatinib in HCC1954 cells, the total amount of GSDMB protein was not clearly increased in gastroesophageal cancer cells (OE19 and N87) neither after lapatinib nor trastuzumab treatment, since in these cells the appearance of a processed form of GSDMB protein (p37) was evident (**Fig. 1B** and Supplementary Fig. S2B). This protein fragment, detected at the latest treatment time points, corresponds to the previously identified C-terminal cleavage product generated by apoptotic caspase-3 (29) (Supplementary Fig. S2C). GSDMB processing by caspase-3 generates N- (p10) and C-terminal (p37) fragments (29) that do not have effect on cell death induction (17). In fact, this processing is analogous to the caspase-3/7 cleavage of GSDMD that eliminates GSDMD pyroptotic effect on cells undergoing apoptosis (30).

**Figure 1.**
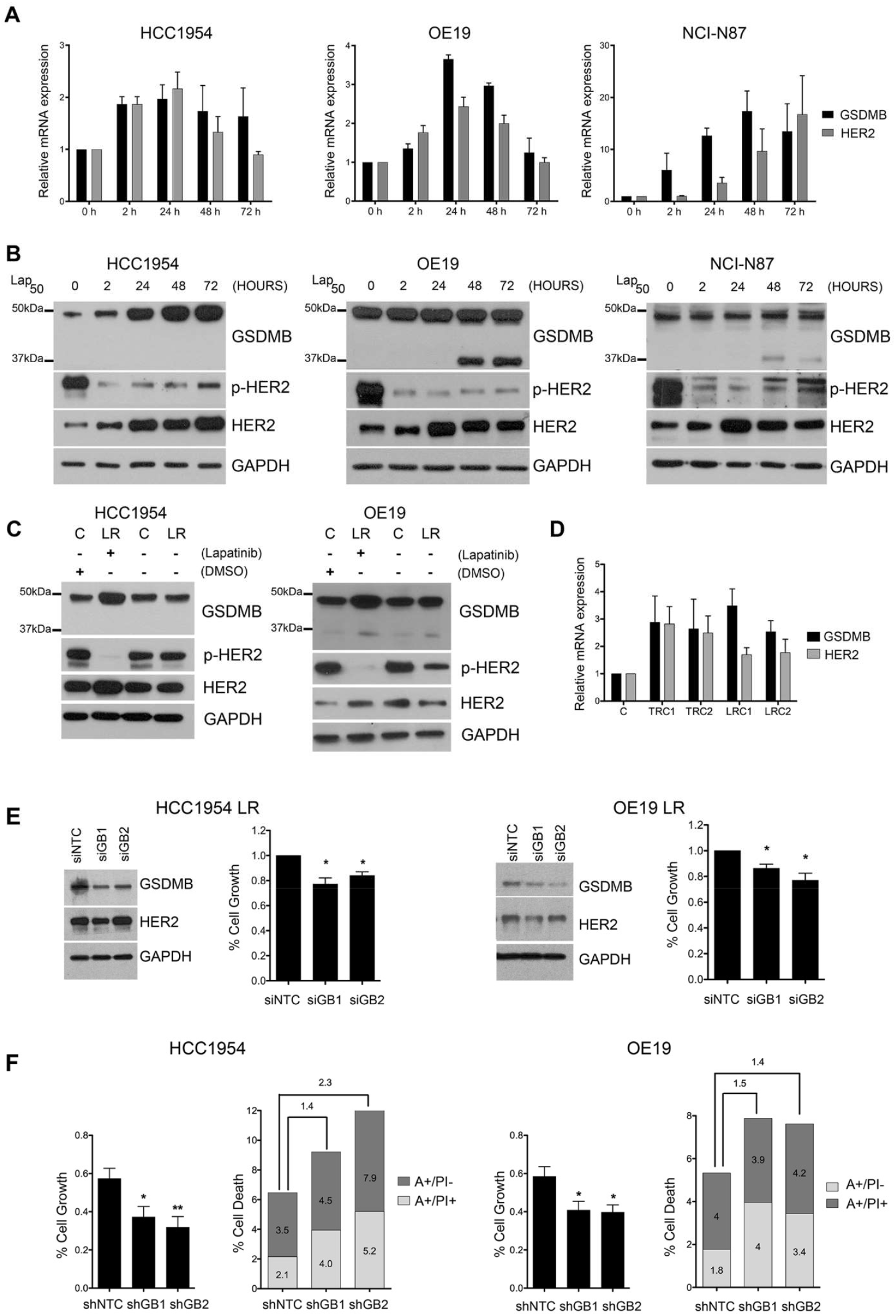
GSDMB is induced in response to anti-HER2 therapies while GSDMB silencing increases the sensitive to lapatinib treatment. **A** and **B**, Relative mRNA (**A**) and protein levels (**B**) of GSDMB and HER2 in HCC1954, OE19 and NCI-N87 cells treated with IC50 of lapatinib (2 µM, 0.7 µM and 0.15 µM, respectively) at different time points. **C**, GSDMB and HER2 protein levels in lapatinib resistant HCC1954 and OE19 cells (LR) and their corresponding control cells (C) treated chronically with lapatinib and DMSO, respectively, and after ten days of drug removal. Study of the different cytotoxic effect of the chronic lapatinib treatment in these cells was analyzed by cell viability assays. **D**, Relative mRNA levels of *GSDMB* and *HER2* in trastuzumab (TRC1 and TRC2) and lapatinib (LRC1 and LRC1) resistant tumors, compared to the parental tumor (C) derived from a HER2 breast cancer Patient Derived Xenograft. **E**, GSDMB expression was reduced in HCC1954 LR (left) and OE19 LR cells (right) by two specific siRNAs (siGB1 and siGB2), in comparison with the control (siNTC). **F**, Cytotoxic effect after 72 h treatment with IC50 of lapatinib (2 µM and 0.7 µM, respectively) in GSDMB-shRNAs-silenced cells HCC1954 and OE19 (right) cells was assessed by cell growth (viability assays, left panel) and death (Annexin V FITC and PI, right panel). Annexin V-FITC positive cells alone (A+/PI-) and Annexin V-FITC and PI doubled stained (A+/PI+) were defined as apoptotic cells. The number over the bars indicate the ratio of cell death relative to the shNTC condition. Statistical significance was determined by two-tailed unpaired *t*-test (^*^*P* < 0.05; ^**^*P* < 0.01). Data are shown as the mean ± s.e.m. Three independent experiments with similar results were performed. In (**A** and **D**), gene expression was normalized to the mRNA levels of *GAPDH*. A, Annexin V; PI, propidium iodide. NTC, non-targeting control. DMSO, dimethyl sulfoxide.

Next, to assess further if GSDMB induction correlates with acquired resistance to HER2-targeted therapies, we generated HCC1954 and OE19 cells long-term resistant (LR) to high doses (>IC50) of lapatinib. In both models, again we observed an upregulation of GSDMB and HER2, and as expected a strong decrease in HER2 phosphorylation (28), compared to their respective control cells (chronically grown with high doses of the vehicle DMSO) (**Fig. 1C**). Interestingly, GSDMB upregulation was dependent on lapatinib presence, since drug removal for 10 days restored GSDMB protein levels (**Fig. 1C**). Moreover, as a complementary model, we used primary cultures from a previously generated HER2+ breast cancer Patient Derived Xenografts (26). Thereby, compared to the parental cell line (C), which is sensitive to trastuzumab, the resistant clones to trastuzumab (TRC1 and TRC2) or lapatinib (LRC1 and LRC2) showed a sharp GSDMB upregulation (**Fig. 1D**).

All these data together with our published observations in PDXs and HER2 breast carcinoma patients (6) reinforce the association between GSDMB over-expression and resistance to different anti-HER2 therapies not only in breast but also in gastric tumors.

### GSDMB increases pro-survival autophagy in response to lapatinib treatment

In order to study if GSDMB is functionally involved in the early response and long-term resistance to lapatinib, we silenced GSDMB in OE19 and HCC1954 cells by shRNA (two different sequences shGB1 and shGB2), as previously reported (7,8). These shRNAs robustly decrease GSDMB expression at the protein (Supplementary Fig. S2D) and mRNA levels (all four GSDMB isoforms that translate to protein) in OE19 (Supplementary Fig. S2E) and HCC1954 cells (8), and do not target other GSDM genes (Supplementary Fig. S2F). Furthermore, in the resistant HCC1954 LR and OE19 LR cells, GSDMB expression was transiently reduced by two different GSDMB-specific siRNAs (siGB1/2, **Fig. 1E**). Stable GSDMB-silencing by lentiviral shRNA transduction could not be obtained in these models because they exhibited intrinsic resistance to the selection antibiotic puromycin.

GSDMB-shRNA-silenced HCC1954 and OE19 cells were significantly more sensitive to lapatinib treatment, as they exhibited an important reduction in cell viability and an increase in cell death compared to shNTC control cells (**Fig. 1F**). Lapatinib treatment did not consistently affect the mRNA levels of other GSDM genes in both cell lines (Supplementary Fig. S2F). Likewise, in the LR models with siGSDMB, we found a slight but statistically significant decrease (around 20%) in cell viability in the presence of lapatinib (**Fig. 1E**). Additionally, GSDMB silencing did not affect the levels of HER2 receptor in any of our cell models (Supplementary Fig. S3A), suggesting that GSDMB does not promote response/resistance to lapatinib through direct modulation of HER2 quantity. To decipher the molecular mechanism by which GSDMB modulates lapatinib response we focused on autophagy, as this process has been demonstrated to act as a resistance mechanism to anti-HER2 therapies both *in vivo* and *in vitro* (22). Autophagy induction during tumor progression can lead to either survival (pro-tumor) or cell death (anti-tumor) depending on the stimulus and the cellular context (24). Hence, we first tested in HCC1954 and OE19 parental cells if lapatinib treatment induced autophagy with survival or death consequences (Supplementary Fig. S3B-S3E). In both cell lines, lapatinib treatment induced autophagic response at 24-48 h, measured by the increase in the levels of LC3B-II (Supplementary Fig. S3B). Importantly, this autophagy is protective since blocking autophagic flux with chloroquine (CQ), which affects the completion of the latter stages of autophagy (31), significantly increased the cytotoxicity of lapatinib (Supplementary Fig. S3C). Similarly, blocking the formation of autophagosomes (32) by an ATG5-specific siRNA (Supplementary Fig. S3D) enhanced the effect on cell viability of lapatinib (Supplementary Fig. S3E).

Therefore, we next assessed whether cells with high or low GSDMB expression could have different endogenous autophagic responses by measuring autophagic flux as the accumulation of the lipidated LC3B (LC3B-II) form in western blots (33,34). In this regard, we analyzed LC3B-II turnover in the presence and absence of lysosomal degradation using CQ. Therefore, higher LC3-II levels were observed in GSDMB-expressing (shNTC) HCC1954 and OE19 cells in comparison with GSDMB-silenced cells in basal autophagy, (**Fig. 2A** and **B**, see black bars) and this effect was significantly exacerbated upon autophagy activation with lapatinib (**Fig. 2A** and **B**, see grey bars). In the same way, HCC1954 LR and OE19 LR cells, which express high levels of GSDMB, exhibit increased LC3B-II accumulation by western blot (**Fig. 2C** and **E**) and LC3B-II puncta by confocal imaging (**Fig. 2D** and **F**) compared to their respective control cells. Thus, these results indicate that high GSDMB expression somehow increases the intrinsic autophagic response to cell stress. Accordingly, we confirmed that the autophagic flux induced by lapatinib (**Fig. 2A** and **B**), or by serum starvation (Supplementary Fig. S4A and S4B) is reduced in GSDMB-silenced cells compared to control lines. Moreover, in HCC1954 LR cells grown in the presence of high doses of lapatinib, the reduction of GSDMB expression by siRNAs also diminishes the autophagic flux (**Fig. 2G**).

**Figure 2.**
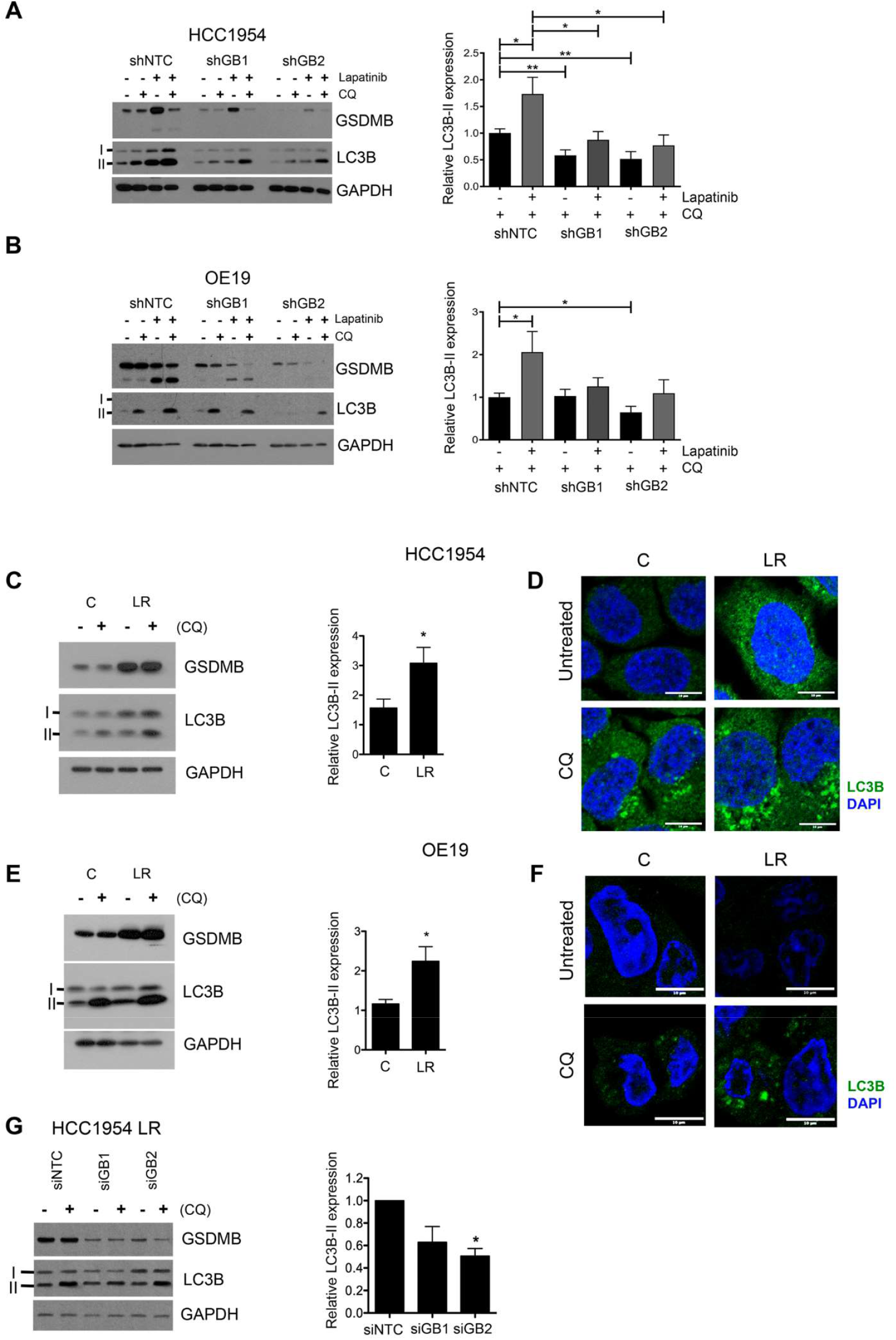
GSDMB-high cells show an increased autophagic flux in response to lapatinib. **A** and **B**, GSDMB and LC3B protein levels in shNTC, shGB1 and shGB2 HCC1954 (**A**) and OE19 (**B**) cells treated with lapatinib (2 µM and 0.7 µM, respectively) and/or chloroquine -CQ- (10 µM and 50 µM, respectively) for 72 h. **C** and **E**, Western blot analysis of GSDMB and LC3B in HCC1954 LR (**C**) and OE19 LR (**E**) cells and their respective controls (C) treated with or without CQ (10 µM and 50 µM, respectively) for 72 h. **D** and **F**, LC3B expression (green) analysis by confocal microscopy in HCC1954 LR (**D**) and OE19 LR (**F**) cells and their controls (C) treated with or without CQ at the concentrations indicated in (**A** and **B**). Representative confocal microscopy images were shown, scale bar, 10 µm. Nuclei were counterstaining with DAPI. **G**, GSDMB and LC3B protein levels in GSDMB-shRNA-silenced HCC1954 LR cells treated with or without 10 µM CQ for 72 h. Quantification of LC3B-II expression (showed on the left of panels, **A-C, E** and **G**) was carried out by densitometric scanning and normalized to GAPDH expression following previous methods (33,34). Statistical significance was determined by two-tailed unpaired *t*-test (^*^*P* < 0.05; ^**^*P* < 0.01). Data are shown as the mean ± s.e.m. Three independent experiments with similar results were performed. NTC, non-targeting control.

Then, given that GSDMB-high cells exhibit increased autophagic lapatinib response, and that this process promotes survival in our cell models (Supplementary Fig. S3), we postulated that GSDMB-high cells could be particularly sensitive to the combination of lapatinib with autophagy inhibitors. Indeed, the autophagy blockage with CQ in GSDMB-high (shNTC) HCC1954 and OE19 cells treated with lapatinib produces a significant reduction on the cell viability (**Fig. 3A** and **C**) as well as an increase in cell death rates (**Fig. 3B** and **D**) compared to lapatinib alone. Nevertheless, no such dramatic effect was observed in the GSDMB-silenced cells. Importantly, in HCC1954 LR and OE19 LR cells, which over-express GSDMB, the addition of CQ strongly increases the cytotoxic effect of lapatinib, and thus revert their resistance to this anti-HER2 drug (**Fig. 3E-H**). Consistent with these results, the autophagy blockage by siATG5 confirmed the increased sensitivity to lapatinib in GSDMB-high cells in all cell models (Supplementary Fig. S5). It should be noted that particularly in OE19 LR cells, which show the highest GSDMB levels, the autophagy inhibition alone (either by CQ or siATG5) has a significant effect on cell viability (**Fig. 3G** and **H** and Supplementary Fig. S5D), supporting that after long-term challenge with this anti-HER2 therapy the subsistence of these cells mostly relies on the endogenous pro-survival autophagic process.

**Figure 3.**
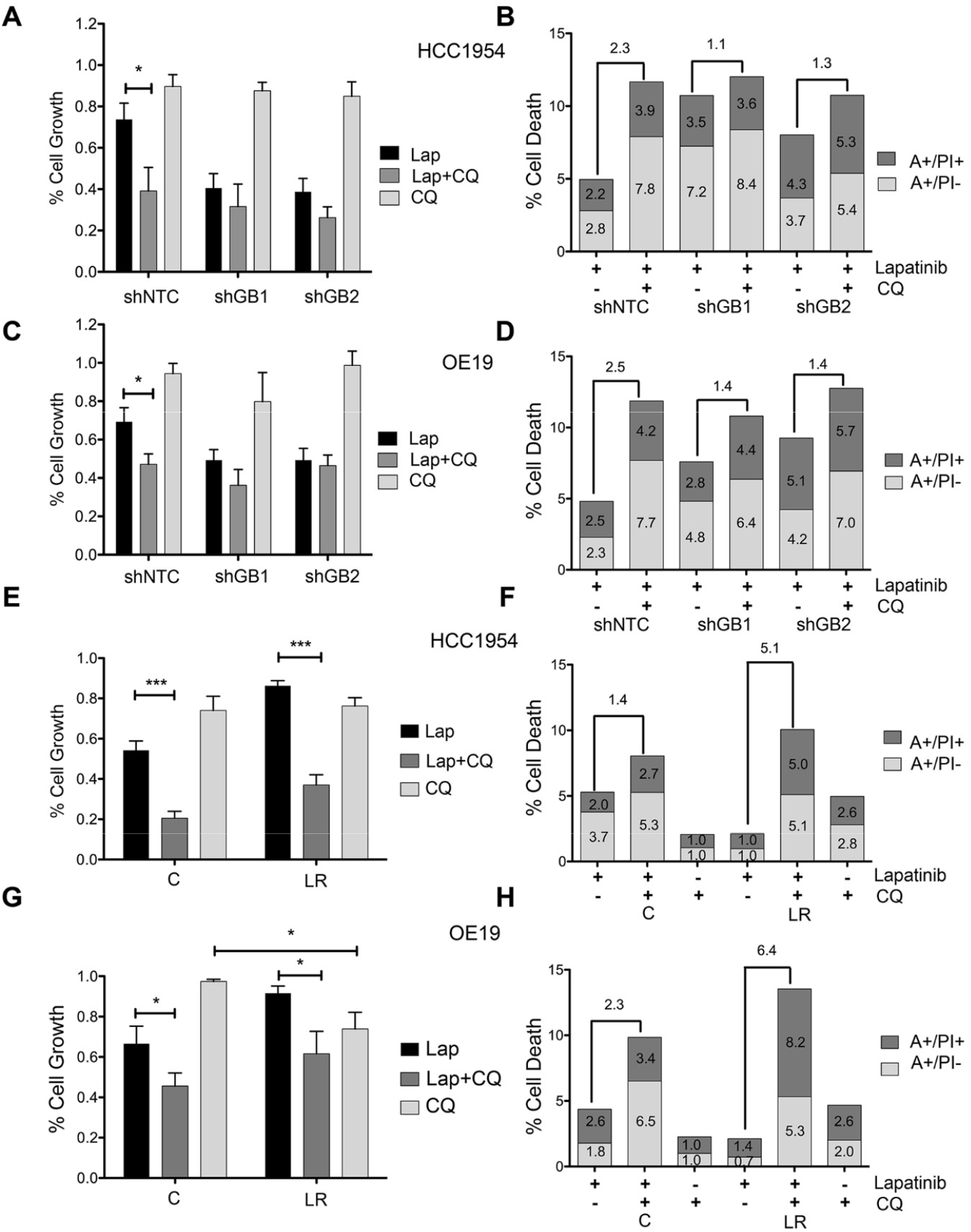
GSDMB-high cells are particularly sensitive to the combination of lapatinib plus chloroquine. **A-D**, the cytotoxic effect of the treatment with lapatinib and/or chloroquine in shNTC, shGB1 and shGB2 HCC1954 (**A** and **B**) and OE19 (**C** and **D**) cells was evaluated by means of cell viability assays and Annexin V-FITC plus PI. **E-H**, The outcome in terms of cell viability and apoptosis of the different treatment regimens was analyzed in HCC1954 LR (**E** and **F**) and OE19 LR (**G** and **H**) cells compared to their corresponding control cells (C). Statistical significance was determined by two-tailed unpaired *t*-test (^*^*P* < 0.05; ^**^*P* < 0.01; ^***^*P* < 0.001). Data are shown as the mean ± s.e.m. Annexin V-FITC positive cells alone (A+/PI-) and Annexin V-FITC and PI doubled stained (A+/PI+) were defined as apoptotic cells. The number over the bars indicate the fold increase in cell death between the indicated conditions (**B, D, F** and **H**). Three independent experiments with similar results were performed. A, Annexin V; PI, propidium iodide; CQ, chloroquine. NTC, non-targeting control.

Taken together, these results confirm that GSDMB enhances the pro-survival autophagy in response to lapatinib in HER2+ breast and gastric cancer cells, and thus GSDMB-over-expressing cells are more sensitive to the addition of autophagy inhibitors both in the early response to HER2-targeted treatment and in drug resistant cells.

### Autophagy inhibition enhances lapatinib efficacy *in vivo* specifically in GSDMB-expressing breast cancer cells

Next, in order to validate if the combination of lapatinib and CQ on HER2/GSDMB positive tumors would be an effective therapeutic approach, we assayed its functional effect on tumor growth *in vivo* using two different preclinical models, zebrafish and mice. In zebrafish, which allows testing the drug response in a large number of biological replicates (35), we first calculated the acute dose-dependent toxicity of both lapatinib and CQ by analyzing the embryo mortality (Supplementary Table S2). Next, the different HCC1954 models mentioned above (shNTC, shGB1/2 as well as LR and the corresponding control cells) that stably express GFP were inoculated into the yolk sac of zebrafish embryos and treated with either lapatinib, CQ or the combination of both (**Fig. 4**). As expected, lapatinib treatment produced a significant reduction in cancer cell growth of GSDMB-silencing tumors (shGB1 and shGB2), but not in HCC1954 shNTC (**Fig. 4A** and **B**). Remarkably, the combination of lapatinib plus CQ provoked a synergistic effect on tumor growth reduction (p<0.001) only in GSDMB-expressing (shNTC) tumors, compared to lapatinib alone, while no such effect was observed in GSDMB-silenced (shGB1, shGB2) tumors. Furthermore, similar results were found on lapatinib resistant cells (LR), where the combined treatment (CQ plus lapatinib) practically abolished the resistant phenotype of HCC1954 LR cells (**Fig. 4C** and **D**). These findings support that the combination therapy was effective specifically in high GSDMB-expressing tumors.

**Figure 4.**
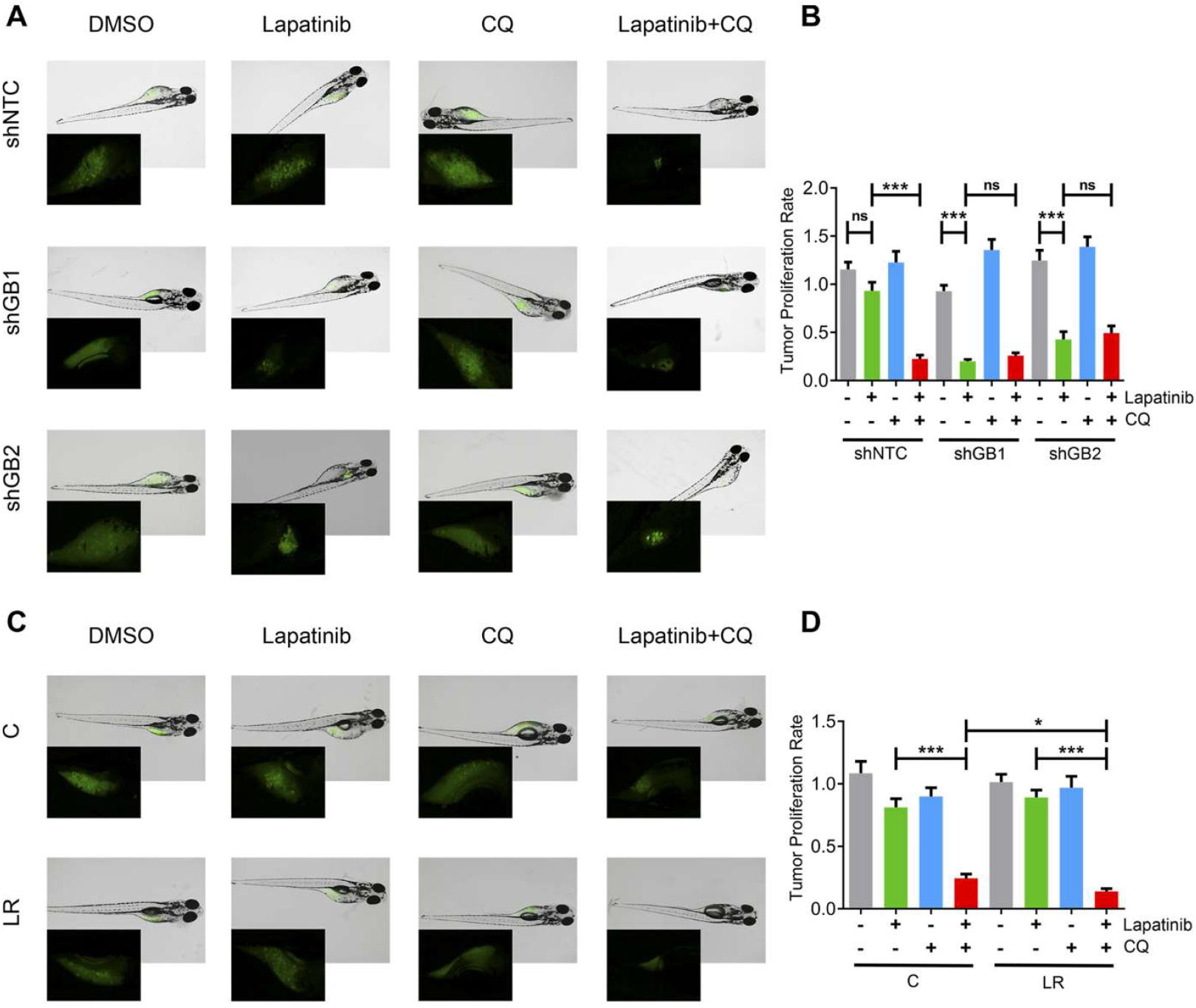
Autophagy blockade with chloroquine improves lapatinib efficacy *in vivo* in zebrafish xenografts of GSDMB-expressing tumors. **A** and **C**, Representative fluorescence stereomicroscope images of GFP expressing control (shNTC), and GSDMB-silenced (shGB1 and shGB2) HCC1954 xenografts (**A**) or HCC1954 LR and control (C) tumors (**C**) treated with the LC50 of lapatinib and/or chloroquine (35,1 mM and 116,4 mM, respectively). Insets represent an augmented image of GFP-positive tumor. **B** and **D**, Tumor proliferation rate of the different HCC1954 xenografts was analyzed by measuring the fluorescence intensity ratio (48 hpt/0 hpt). Statistical significance was determined by two-tailed unpaired *t*-test (^*^*P* < 0.05; ^**^*P* < 0.01; ^***^*P* < 0.001; ns, nonsignificant). Data are shown as the mean ± s.e.m. At least, n = 20 per each indicated condition. CQ, chloroquine; LC50, 50% lethal concentration; hpt, hours post-treatment. NTC, non-targeting control.

Subsequently, to validate these results we performed similar treatment experiments in mice bearing orthotopically injected HCC1954 (shNTC, shGB1 and shGB2) breast cancer xenografts (**Fig. 5** and Supplementary Fig. S6). Once again, while lapatinib alone only decreased significantly tumor growth in GSDMB-silenced tumors (**Fig. 5C** and **D**), the combination of CQ and lapatinib caused a synergistic effect on the reduction of tumor volume exclusively in GSDMB-expressing (shNTC) tumors (**Fig. 5B**). Importantly, no proliferation changes by PCNA immunohistochemical expression were detected in any of the experimental conditions (Supplementary Fig. S6A). As observed *in vitro*, GSDMB was also up-regulated *in vivo* in GSDMB-expressing tumors (shNTC) by lapatinib (**Fig. 5E** and Supplementary S6B), and again it was associated to a reduced response to the treatment compared to GSDMB-silenced tumors (**Fig. 5E**). These *in vivo* data prove that the autophagy blockage is essential to improve the antiHER2 therapy response in HER2/GSDMB tumors and reinforce the role of GSDMB in the promotion of pro-survival autophagy as a resistance mechanism to these therapies.

**Figure 5.**
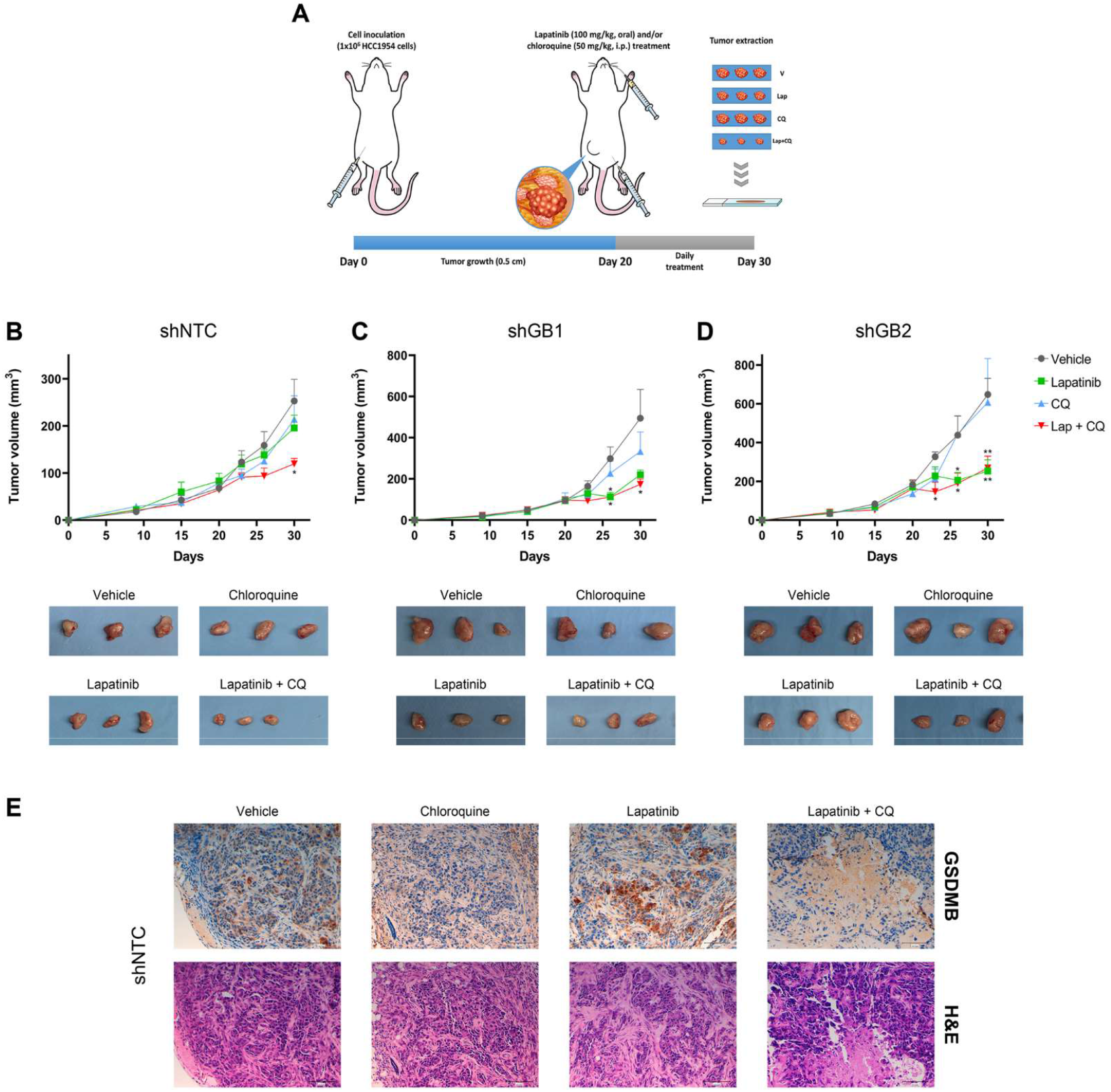
The combination of lapatinib with chloroquine increases the therapeutic response *in vivo* in GSDMB-expressing orthotopic mouse tumor xenografts. **A**, Experimental design of the mouse xenograft model (n = 5 per condition). Mice were inoculated with either HCC1954-mCherry-luc control (shNTC), or GSDMB-silenced cells (shGB1 and shGB2) and treated with lapatinib (100 mg/kg, orally, once daily), CQ (50 mg/kg, intraperitoneally, once daily), or a combination of both (lapatinib+CQ). An aqueous solution containing 0.1% Tween 80 and 0.5% Hypromellose was used as vehicle. **B-D**, Quantification of tumor volume evolution of shNTC (**B**), shGB1 (**C**) and shGB2 (**D**) HCC1954 xenografts, treated with the indicated treatment regimens. Representative images of the tumor size *ex vivo* after each treatment. Statistical significance was determined by multiple unpaired *t*-test – comparing vehicle with each of the other conditions in every time point (^*^*P* < 0.05; ^**^*P* < 0.01). Data are shown as the mean ± s.e.m. **E**, Representative images of GSDMB immunohistochemical analysis and hematoxylin and eosin staining in shNTC tumors, treated with the different therapeutic strategies indicated in (**A**) (scale bar, 100 µm). CQ, chloroquine. NTC, non-targeting control.

### GSDMB and LC3B cooperate in the pro-survival autophagy induction in response to HER2 targeted therapy

Next, to validate the association of GSDMB expression and pro-survival autophagy in clinical samples, we studied the immunohistochemical expression of LC3B, as a marker of autophagy status, in a cohort of HER2 breast carcinomas in which the GSDMB over-expression/amplification was previously reported (6) (**Fig. 6A**) as well as in a novel series of HER2 gastric tumors (Supplementary Fig. S1A and S1B). As reported before (36), a strong LC3B puncta staining suggests autophagy induction. In this regard, about 50% of breast and gastric tumors showed intense L3CB dotted expression (**Fig. 6A**, Supplementary Fig. S1A, Supplementary Table S1 and S3). In this sense, strong LC3B puncta was more frequent in GSDMB over-expressing breast and gastric tumors (11/18 cases, p=0.007, and 10/15 p=0.045, respectively; **Fig. 6A** and Supplementary Fig. S1B). Moreover, although GSDMB staining is mainly diffuse cytoplasmic or nuclear (6,8,37) a similar dotted co-expression was observed for GSDMB and LC3B puncta within tumor cells (**Fig. 6A**, Supplementary Fig. S1A and S1B). Importantly, the GSDMB/LC3B puncta positive co-expression strongly correlated with relapse in GSDMB+ breast and gastric carcinomas (8/11 p=0.001, 9/10 p=0.024, respectively; **Fig. 6A**). These findings support that those tumors with GSDMB over-expression, which have poor prognosis, could exhibit intrinsic autophagy, even in clinical specimens.

**Figure 6.**
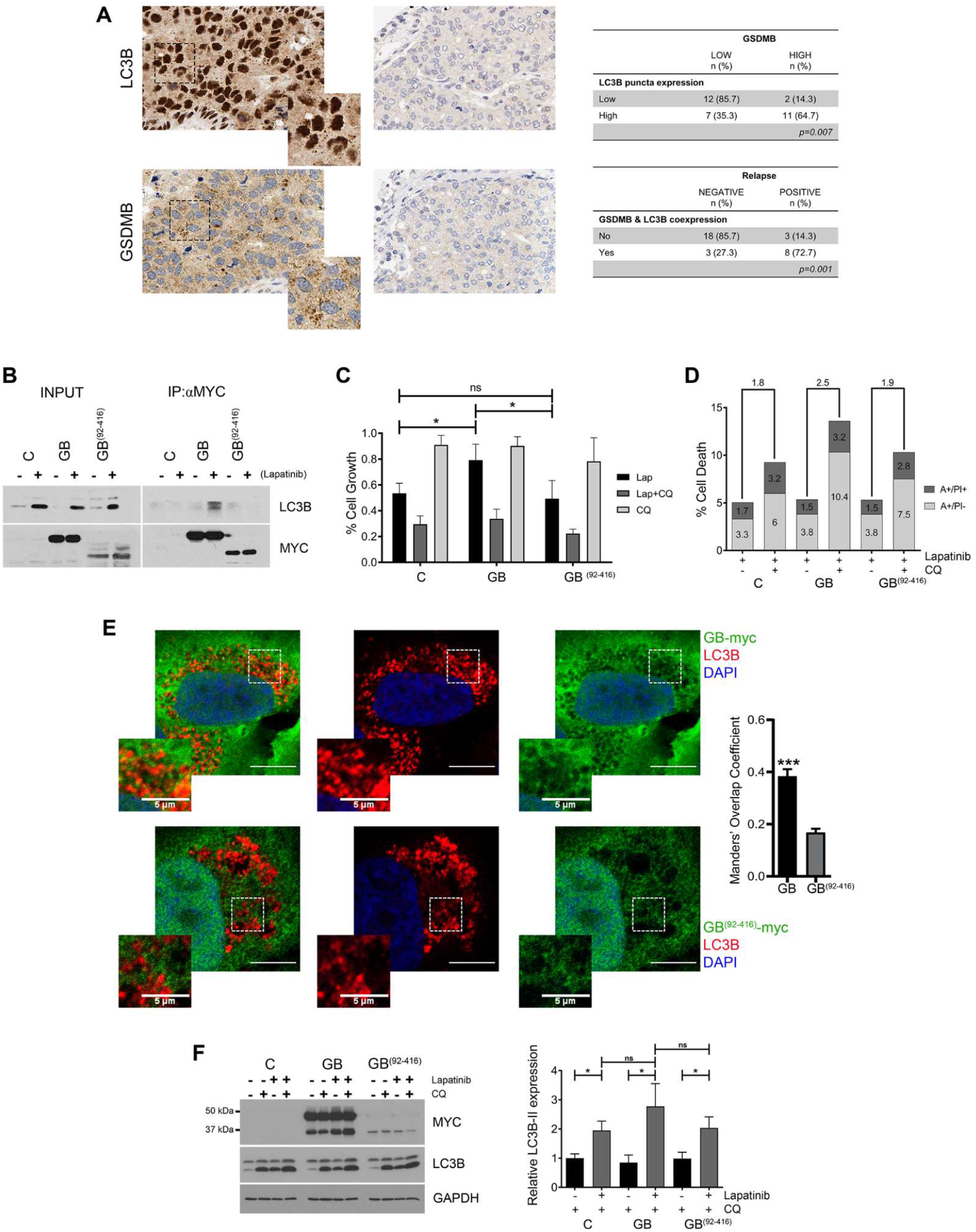
GSDMB modulates autophagic response through LC3B interaction. **A**, Representative immunohistochemistry images of LC3B and GSDMB positive (left) and negative (right) expression in a cohort of adjuvant treated HER2 breast carcinomas (6). Insets are magnified images of the boxed areas (scale bar, 10 µm), highlighting the similar localization of GSDMB and LC3B puncta staining. Correlation between GSDMB and LC3B expression as well as GSDMB/LC3B co-expression with relapse in HER2 breast carcinomas (right). Differences were compared by Fisher’s exact test. **B**, Co-immunoprecipitation assay of LC3B-GSDMB interaction after autophagy induction with lapatinib in HCC1954 cells exogenously expressing full length GSDMB (GB), a construct lacking the LIR motifs (GB^(92-416)^) or control (C; empty vector) cells. **C** and **D**, Comparative cytotoxic effect by cell viability assays (**C**) and Annexin V-FITC plus PI (**D**) in HCC1954 C, GB and GB^(92-416)^ cells after 72 h treatment with lapatinib (2 µM) and/or CQ (10 µM). Annexin V-FITC positive cells alone (A+/PI-) and Annexin V-FITC and PI doubled stained (A+/PI+) were defined as apoptotic cells. **E**, Representative images of the colocalization between LC3B (red) and GSDMB-Myc-tag (green) by confocal microscopy in HCC1954 GB and GB^(92-416)^ cells after lapatinib (2 µM) plus CQ (10 µM) treatment. Nuclei were counterstaining with DAPI. Quantification of the Manders’ Overlap Coefficient (LC3B overlapping Myc-tag) is shown on the right. Five independent experiments were performed obtaining a total of 70 cells, per experimental condition. Scale bar, 10 µm. **F**, Western blot analysis of Myc-tagged GSDMB and LC3B in HCC1954 C, GB and GB^(92-416)^ treated for 72 h with lapatinib (2 µM) and/or CQ (10 µM). Quantification of LC3B-II expression was performed by densitometric scanning and normalized to GAPDH expression. Statistical significance was determined by two-tailed unpaired *t*-test (^*^*P* < 0.05; ^**^*P* < 0.01; ^***^*P* < 0.001; ns, nonsignificant). Data are shown as the mean ± s.e.m. In (**C-E** and **G**), three independent experiments with similar results were performed.

Finally, to decipher whether there is a functional cooperation between GSDMB and LC3B in the autophagy induction, we first analyzed if GSDMB protein contains putative LC3B-interacting region (LIR) motifs using the iLIR database (38). Two putative LIR motifs were detected on the GSDMB N-terminal domain, “LIR1” corresponding to ^3^SVFEEI^8^ sequence and “LIR2” covering ^82^AEFQIL^87^ aminoacids (Supplementary Fig. S7A). In parallel, we carried out a protein interaction prediction assay of GSDMB and LC3B using the algorithm HawkDock (39). This study showed an energetically feasible (−2 Kcal/mol) GSDMB/LC3B protein-protein complex, in which the LIR1 domain of GSDMB and the N-terminal domain of LC3B (^38^YKGEKQ^43^) are potentially implicated (Supplementary Fig. S7B). To investigate if these LIR motifs contribute to GSDMB function in autophagy regulation we generated HCC1954 cells exogenously over-expressing either myc-tagged full length GSDMB (GB, **Fig. 6B**) or a deletion construct lacking both LIR motifs (GB^92-416^, which corresponds to the C-terminal fragment released by caspase-3 processing (29). Cells stably expressing those constructs were treated with lapatinib to induce autophagy and then we carried out immunoprecipitation assays using anti-myc (**Fig. 6B**) and anti-LC3B antibodies (Supplementary Fig. S7C). Remarkably, LC3B was only immunoprecipitated in HCC1954 GB cells and not in GB^92-416^ or control conditions (**Fig. 6B**). Furthermore, only GB-myc was immunoprecipitated using anti-LC3B antibody (Supplementary Fig. S7C). In addition, we observed by confocal microscopy that the colocalization of L3CB and GB-myc was significantly higher than LC3B with GB^92-416^ (**Fig. 6E**). Taking together, these data suggest that the GSDMB N-terminal region comprising the aminoacids 1-91 (which contains both putative LIR motifs) is necessary for the interaction with LC3B and may participate in the autophagy induction after lapatinib. In fact, in response to lapatinib a significant cell growth increase (**Fig. 6C**) was observed in GB over-expressing cells in comparison to control cells (expressing GSDMB endogenous levels) or GB^92-416^ cells (**Fig. 6C**). As expected, after the combination of Lapatinib plus CQ the cell death rate was higher in GB expressing cells than the other conditions. Again, control and GB^92-416^ cells showed a similar phenotype indicating that the construct lacking the GSDMB construct lacking the LIR domains is unable to affect the autophagy response (**Fig. 6D**). Furthermore, although the basal autophagy induced by CQ treatment in GB, GB^92-416^ and control HCC1954 cells was quite similar, regarding LC3-II increasing, the increase in autophagic flux after lapatinib plus CQ treatment was stronger in GB cells, regarding LC3-II increasing, compared to the other cell models (grey bars, **Fig. 6F**). Lastly, we verified that GSDMB, is not accumulated after autophagy flux blockage by CQ, in contrast to LC3B or p62, supporting that GSDMB does not behave like an autophagy cargo (40) (Supplementary Fig. S7D).

Taken together, all the above-mentioned results reveal that the autophagy inhibition reverses, at least in part, the resistance to lapatinib mediated by GSDMB over-expression in HER2 cancer cells. Therefore, the combination of autophagy inhibitors with anti-HER2 agents could provide new therapeutic options for GSDMB-overexpressing HER2+ breast and gastroesophageal tumors (Supplementary Fig. S7E).

## DISCUSSION

GSDMB arises as a new biomarker in cancer (10) although plays a complex role in tumor biology. On one hand, its upregulation occurs in diverse cancer types (6–8,10,37), where usually associates with poor prognosis (6– 8,37). In particular, in HER2 breast tumors the GSDMB co-amplification/over-expression (occurring in >60% of cases) promotes an aggressive behavior by potentiating cell invasion, metastasis and drug resistance (6–8). On the other hand, similar to other GSDM family members (10,12,14), it possesses a potentially “activatable” pro-cell death function (8,16,19), that could be possibly exploited as a therapeutic approach in GSDMB-overexpressing cancers. In fact, our previous work proved that GSDMB pro-tumorigenic actions could be reverted partially *in vitro* and *in vivo* by a GSDMB-antibody-based nanomedicine that seemed to activate the protein intrinsic cytotoxic activity (8). Furthermore, a recent study evidenced that the cytotoxic activity of GSDMB can be released in tumor cells upon proteolytic cleavage of its N-terminus by lymphocyte Granzyme-A (19). However, triggering *in vivo* GSDMB-mediated tumor pyroptosis required pharmacological activation of the antitumor immunity to achieve partial tumor regression (19). Despite promising results, these approaches based on GSDMB cytotoxic function require further development in order to get into the clinic.

In the present work, we have focused in tackling specifically the mechanisms by which GSDMB mediates limited response to anti HER2 therapy. The data presented with lapatinib, combined with our previous results with trastuzumab (6), prove that GSDMB is specifically upregulated (mRNA and protein levels) in response (early response and during acquired resistance) to different anti-HER2 targeted drugs in HER2 breast cancer cell lines and PDXs, as well as gastroesophageal tumor cells. Moreover, GSDMB silencing by si-/shRNAs increases sensitivity to these drugs thus reverting partially the resistant phenotype.

Interestingly, here we have proven for the first time that GSDMB-mediated therapy resistance requires activation of protective autophagy. Autophagy is a well-described catabolic mechanism where molecules and damaged cellular organelles are degraded (41) and it has been implicated in the etiology of cancer, acting as either a pro-death or a pro-survival factor depending on the type and stage of cancer (41). In this regard, both lapatinib (22) and trastuzumab (42) have been shown to trigger protective autophagy as a resistance mechanism whereby tumor cells display autophagy addiction to maintain the resistant phenotype (43). Accordingly, abrogation of autophagy by specific inhibitors like chloroquine (CQ), which affects the completion of the latter stages of autophagy (31), can re-sensitize cancer cells to these drugs, and in fact, preclinical studies suggest that CQ in combination with HER2-targeted inhibitors enhance tumor cell death (43). Nevertheless, the biological determinants affecting autophagic response to these drugs and the sensitivity to autophagy blocking agents are not well understood so far.

Here, we uncover a novel function of GSDMB as a modulator of the autophagy response to anti-HER2 therapies in breast and gastric cancers. First we show that GSDMB-high cells exhibit higher levels of LC3B-II, the lipidated form of the autophagy marker LC3B necessary for autophagosome binding (44), upon lapatinib and CQ co-treatment, than in GSDMB knockdown cells. Moreover, in archived breast tumors specimens (where GSDMB-driven resistance to trastuzumab has been reported (6) and gastric tumors we observed that LC3B protein puncta expression, denoting LC3B-II, was globally higher in HER2-positive tumors. In line with previous reports describing GSDMB (6,8) and LC3 over-expression (45,46) as cancer risk factors, we confirmed that breast and gastric tumors which high puncta expression of both GSDMB and LC3B are prone to relapse, thus reinforcing the link between GSDMB expression, active autophagy flux and poor prognosis. Notably, in diverse cellular models the inhibition of autophagy flux with CQ significantly enhances responses to lapatinib *in vitro*, inducing a shift from cell viability towards toxicity both in early response and in lapatinib-resistant GSDMB-expressing cells. Combination of lapatinib and CQ nearly abrogates tumor growth in preclinical *in vivo* (zebrafish, mouse) models, demonstrating the potential for a synergistic therapeutic strategy to reverse the HER2-targeted drug resistance effects of GSDMB.

It should be pointed out that other GSDM members have been associated with autophagic processes, but sometimes this function requires GSDM processing by proteases (47). For instance, in neutrophils, caspase-1 cleaved GSDMD N-terminal domain, but not full length GSDMD, localizes to the membrane of LC3+ autophagosomes, inducing autophagy-mediated IL-1b secretion independent of pyroptotic activity (48). Moreover, GSDMA3 mutant proteins lacking auto-inhibition or the free GSDMA3 N-terminal region can increase LC3-II and autophagy in HEK293 cells, though in this case, this leads to cell death (15). In contrast, normal full length PJVK (DFNB59), recruits LC3-II to produce pexophagy (49); a type of selective autophagy for degrading damaged peroxisomes (50). Our data revealed protein-protein interaction between the N-terminal domains of LC3B and GSDMB; possibly through specific putative LC3B-interacting motifs (LIR, ^3^SVFEEI^8^ and ^82^AEFQIL^87^) located on the GSDMB N-terminus. Indeed, full length (uncleaved) GSDMB binds to LC3-II and may facilitate activation of protective autophagic responses to lapatinib. However, during lapatinib induced cell death, the activation of pro-apoptotic caspases 3/7 partially cleave GSDMB within its N-term (^88^DNVD^91^) to generate two fragments. Alike the analogous processing in GSDMD (30), these products do not have intrinsic effect on cell death (17). Moreover, we showed that the C-terminal GSDMB cleaved fragment (GB^92-416^, lacking LIR motifs) does not bind to LC3 nor does it affect the autophagy response. Whether the GSDMB N-term fragment (containing the LIR motifs) retains the activity to bind LC3B is still unknown.

Even though it remains unclear how the N-terminal domain of GSDMB interaction with LC3B modulates autophagy, we postulate that this region might contribute to LC3-II anchorage to autophagosomes, since both the N-terminus in GSDMB and the full protein bind specific lipids such as sulfatides and, more weakly, cardiolipin (8,29). Hence, during the initial stages of autophagy induction LC3 would be rapidly cleaved to produce cytoplasmic LC3-I, followed by lipidation of the latter into LC3-II (51), preferably with phosphatidylethanolamine, cardiolipin and other dianionic lipids (52,53). Such events may thus enable recruitment of LC3-interacting proteins through LIR domains for the growth of autophagic membranes (44).

In summary, we have established that GSDMB-mediated resistance to anti-HER2 therapy may result from induction of pro-survival autophagy. We also provide evidence for the utility of combination therapies based on the autophagy inhibitor CQ and the HER2-targeted drug lapatinib; thus, contributing to the evolving treatment paradigm in HER2 tumors overexpressing GSDMB, which largely associate with poor outcomes.

