## Supplementary data for "Gasdermin B over-expression arbitrates HER2-targeted therapy resistance by inducing protective autophagy" (from another version of this preprint: version 1, 2021-07-01)

**Supplementary Methods.**

**Supplementary Tables S1-5.**

**Supplementary Table S1.** Summary of the clinic-pathological data of the gastric carcinoma cohort.

**Supplementary Table S2.** *In vivo* acute toxicity results in zebrafish.

**Supplementary Table S3.** Summary of the clinic-pathological data of the adjuvant treated HER2 breast carcinoma series.

**Supplementary Table S4.** List of primers used for qRT-PCR and cloning.

**Supplementary Table S5.** List of primary antibodies used for western blot (WB), immunofluorescence (IF), immunoprecipitation (IP), and immunohistochemistry (IHC).

**Supplementary Figures S1-7.**

### SUPPLEMENTAL METHODS

**RNA interference and overexpression constructs.** GSDMB-silenced HCC1954 and OE19 cells were generated using Mission® shRNA Lentiviral Transduction Particles (SHCLNV-NM_018530, Sigma-Aldrich, shGB1: TRCN0000137108 & shGB2: TRCN0000168794) according to the manufacturer's instructions. Non-targeting Control shRNA Transduction Particles (SHC002V, Sigma-Aldrich) were employed as control (shNTC). Stably infected cells were selected using 0.5 µg/ml puromycin. For HCC1954 LR and OE19 LR cells, GSDMB transient silencing was performed by two specific siRNAs (Custom siRNA, Dharmacon, siGB1: Sense sequence, 5'-GCCAAAGGGAAGUGACCAUUU-3'; Antisense sequence, 5'-AUGGUCACUUCCCUUUGGCUU-3' & siGB2: Sense sequence, 5'-GAGAUUACCCAAAGAAAUAUU-3'; Antisense sequence: 5'- UAUUUCUUUGGGUAAUCUCUU-3'). ATG5 was silenced by a siGENOME Human ATG5 siRNA – SMARTpool (M-004374-04-0010, Dharmacon). siGENOME Non-targeting siRNA (D-001210-01-20, Dharmacon) was utilized as control (siNTC). Transfections were carried out using DharmaFECT 1 Transfection Reagent (T-2001-02, Dharmacon) according to the manufacturer's protocol. HCC1954 cells overexpressing myc-tagged full-length GSDMB (GB,(1) and a GSDMB construct lacking the residues 1-91(GB$^{92-416}$) were obtained by stable lentiviral infection using the pLVX vector. The coding region of GB$^{92-416}$, tagged with a myc-epitope in the C-terminal domain (highlighted in bold), was amplified by PCR using the primers indicated in the **Supplementary Table 4.** The resulting GB$^{92-416}$-myc cDNA was then cloned into the EcoRI and XbaI sites of the pLVX-Puro plasmid (125839, Addgene). Lentiviral particles were generated in HEK293T by transfection (Lipofectamine™ 2000 Transfection Reagent, Invitrogen) with PAX2, VSVG and pLVX-Puro-GB (92-416) plasmids. Selection of infected HCC1954 cells was achieved with puromycin (0.5 µg/ml).

**Western blot and Co-immunoprecipitation.** Protein extracts from cultured cells were obtained using a standard 2% SDS buffer with protease and phosphatase inhibitors for 30 min at 4ºC. Equal amount of proteins measured using the Pierce™ BCA Protein Assay Kit (Thermo Scientific) were resolved by SDS-PAGE and analyzed by immunoblot. Membranes were incubated with the indicated antibodies: HER2 (2248), p-HER2 (Tyr1221/1222) (2243), Myc-tag (2276), Myc-tag (2278) and SQSTM1/p62 (8025) from Cell Signaling Technology; ATG5 (A0731), LC3B (L7543) and GAPDH (CB1001) from Sigma-Aldrich; Caspase-3 (sc-56053) from Santa Cruz Biotechnology; and GSDMB antibody was generated in the lab (2). Amersham ECL HRP-conjugated anti-mouse or anti-rabbit secondary antibodies were used (NA931 and NA934, GE Healthcare Life Sciences, 1:5000 dilution). Detection was carried out with Pierce™ ECL Western Blotting Substrate (Thermo Scientific) and quantification was carried out by densitometric scanning and normalized to GAPDH expression. For co-immunoprecipitation, 2x10$^{6}$ HCC1954 cells were lysed in a buffer containing 50 mM Tris-HCl (pH 7.5), 150 mM NaCl, 10 mM EDTA, 10 mM EGTA, 0.5% NP-40 in presence of a mixture of proteases inhibitors (cOmplete™, Mini Protease Inhibitor Cocktail, Roche). One mg of protein lysate was incubated overnight at 4 ºC with the Myc-tag (2276, Cell Signaling Technology) or LC3B (sc-376404, Santa Cruz Biotechnology) antibody followed by incubation with Dynabeads™ protein G (Invitrogen) for 3 h at 4°C. After washing the beads with the lysis buffer three times, the beads-protein binding was detected by western blotting.

**Immunofluorescence staining.** Cells were grown to 50-70% confluency on coverslips in 24-well plates. After treatment, cells were fixed with 4% paraformaldehyde for 20 min at room temperature (RT). After 5 min permeabilization with 0.1% Triton™ X-100, coverslips were incubated with corresponding primary antibodies (**Supplementary Table 5)**, Myc-tag (2276, Cell Signaling Technology) or LC3B (L7543, Sigma-Aldrich) followed by incubation with secondary antibodies, Alexa Fluor 488 or 546-labeled (A-11029, A-11034 and A-11035, D1306, Invitrogen) for 1 h at RT. Cell nuclei were stained with 1:5000 DAPI (Invitrogen). Images were captured on a LSM710 confocal microscope (ZEISS), analyzed and quantified using ImageJ software. Colocalization analyses of LC3B and Myc-tag were performed by means of calculating the Manders' Overlap Coefficient (LC3B overlapping Myc-tag), using the JACoP plugin in ImageJ (3).

**Drugs and autophagy analysis.** Trastuzumab (Herceptin, Roche) was directly diluted on cell culture medium with inactivated FBS and used at final concentrations of 1 mg/ml (HCC1954 cells) or 400 µg/ml (OE19 and NCI-N87). Lapatinib (Tykerb, GlaxoSmithKline) was dissolved in DMSO (Merck) to a stock concentration of 2.5 mM and used at 2µM (HCC1954), 0.7µM (OE19) or 0.15µM (NCI-N87). Chloroquine diphosphate salt (CQ, Sigma-Aldrich) was applied at 10µM (HCC1954) or 50µM (OE19). The autophagic flux quantification was performed measuring LC3B-II expression by WB as described before (4,5). Briefly, $15 \times 10^4$ HCC1954 and $60 \times 10^4$ OE19 cells/well were seeded in 6-well plates. After 24 h, cells were treated for 72 h with CQ and/or lapatinib or starvation (complete media with 0.01% FBS) with/without CQ. For the analysis of LC3B by immunofluorescence, HCC1954 and OE19 cells were grown on coverslips to 50-70% confluency and then treated for 24 h with IC50 of lapatinib and/or CQ. LC3B staining was studied by confocal microscopy as described above.

**Cell viability assay.** The cytotoxic effect of analyzed compounds on HCC1954, OE19 and NCI-N87 was tested using the AlamarBlue reagent (Bio-Rad). Briefly, $5\times10^3$ HCC1954, $2\times10^4$ OE19 and NCI-N87 cells/well were seeded into a 96-well plate in triplicates. After 24 h, cells were treated with previously indicated concentrations of specific treatments, alone or in combination, for 72 h. Spectrophotometric analysis was performed at wavelengths of 570 and 600 nm.

**Cell apoptosis analysis.** Apoptotic rate was quantified using the Annexin V-FITC stained apoptosis detection kit (Immunostep) and propidium iodide (PI, Sigma-Aldrich) according to the manufacturers' instructions. HCC1954 and OE19 cells were treated with lapatinib and/or CQ at previously described concentrations for 72 h. Annexin V-FITC positive cells alone (A+/PI-) and Annexin V-FITC and PI doubled stained (A+/PI+) were defined as apoptotic cells. The apoptosis analysis was conducted using a FC500 flow cytometer (Beckman Coulter).

**RNA isolation and RT-qPCR.** Total RNA was extracted from cells with the TRIzol™ Reagent (Invitrogen) and the RNeasy Mini Kit (Qiagen), performed following standard procedures. Total RNA from PDX tissue was isolated using the ReliaPrep™ RNA Miniprep System (Promega), according to the manufacturer's protocol. DNase treatment was performed to avoid the presence of residual DNA. cDNA reverse transcription was carried out with the M-MLV RT (VWR Chemicals). Each PCR reaction was performed by triplicate using the PerfeCTa™ SYBR® Green FastMix™ (Quantabio) in a StepOnePlus™ Real-Time PCR system (Applied Biosystems).

Expression values for each gene were normalized to housekeeping gene GAPDH expression. GSDMB isoforms qPCR was conducted using TaqMan gene expression assay primers (GSDMB-1: Hs00938445_m1; GSDMB-2: Hs00939390_m1; GSDMB-3&4: Hs00940508_m1; GSDMB-4: Hs00940509_m1; and GAPDH: Hs02758991_g1; Thermo Fisher Scientific). The rest of qPCRs was performed utilizing the primer sequences indicated in the Supplementary Table **S4.** RT-qPCR analysis were performed in a StepOnePlus™ Real-Time PCR system (Thermo Fisher Scientific).

**Zebrafish studies.** Wild-type zebrafish (AB strain, *Danio rerio*) were maintained in a controlled aquatic facility at 27ºC (± 1ºC) and 14/10 h light/dark light cycle, according to previously described recommendations (6). First, to determine several toxicological parameters of lapatinib and CQ in zebrafish model, the Fish Embryo Acute Toxicity test (FET, following the recommendation of the Organization for Economic Co-Operation and Development (OECD) No. 236, 2013) was performed. Briefly, 72 hours post-fertilization (hpf) hatched embryos were incubated in 24-well plates with different concentrations of the tested drugs for 96 h. Osmosis water and 0.1% DMSO (Merck) were used as negative controls. Three replicates were performed, analyzing a total amount of 60 embryos per concentration. Treated embryos were checked under an inverted optical microscope (Nikon TMS) at 24, 48, 72 and 96 h, in accordance with the guideline test and following the FET test scheme described previously (7). ToxRat software was used to determine the 10% lethal concentration (LC10), 50% lethal concentration (LC50), lowest observed effect concentration (LOEC) and no-observable effect concentration (NOEC) values of the tested drugs. The LC50 values (35.1 mM and 116.4 mM for lapatinib and CQ, respectively) were higher than *in vitro* approaches (2 µM, lapatinb and 10 mM for CQ) confirming that drug concentrations used for xenograft treatment were appropriate as they were not harmful to the host embryos. Next, to analyze the *in vivo* efficacy of the lapatinib and/or CQ, $1x10^6$ of HCC1954 GSDMB expressing and silencing cells (shGB1/shGB2) as well as HCC1954 LR and C resistant cells (expressing green fluorescent protein, GFP ) were inoculated into the perivitelline space of fish embryos of 48 hpf (n= 40-50 embryos per condition). Previously to injection, embryos were anesthetized with 0.003% tricaine (Sigma-Aldrich) and GFP transduced cells were resuspended in 10 µl 2% Polyvinylpyrrolidone (Sigma-Aldrich). Injected embryos were incubated at 34ºC until the end of the experiments. Xenografts were photographed at 24 hpi (hours post-injection) and before drug treatment (0 hpt) with an AZ-100 fluorescence stereomicroscope (Nikon). Then, embryos were randomly distributed in the treatment groups: control (0.1 % DMSO, vehicle), lapatinib (2 µM) and a combination of lapatinib with CQ (10 µM). After 48 hpt, xenografts were photographed again to analyze the tumor growth. At the end of experiments, embryos were euthanized by tricaine overdose. Three experiments were performed obtaining at least 60 embryos per each condition. Tumor growth and treatment response were analyzed measuring the fluorescence intensity ratio (48 hpt/0 hpt) using an image analysis program (QuantiFish, (8)). These ratio values were: 1, when tumor size was maintained; >1, when tumor cells proliferated during the incubation time with the treatment and <1, when tumor cells died after 48 h of incubation (possible outliers, analyzed with the outlier calculator tool of GraphPad software, were discarded). *In vivo* zebrafish acute toxicity parameters were obtained using the ToxRat software. Lethal concentrations (LC) were determined using a linear maximum likelihood regression from the 96 h survival data and with 95% confidence limits. NOEC and LOEC values were obtained using a qualitative trend analysis by contrast, step-down Cochran-Armitage test and Tarone's test. In accordance with OECD guideline, tests were considered valid whether the zebrafish embryos mortality in the negative control was less than 10% and the mortality in the positive control was more than 30%. The number of samples (n) has been indicated for each figure panel. In all cases, $P$ values < 0.05 were considered as statistically significant.

**Mouse studies.** Orthotropic breast tumor xenografts and experimental lung metastasis assays were performed in female nu/nu mice (Charles River) following standard procedures (9). mCherry-luc transduced shNTC, shGB1 or shGB2 HCC1954 ($1x10^6$) cells were injected into the right fifth mammary fat pad (mfp) of 7-week-old female NMRI-nu Immunodeficient Mice (Janvier Labs) and tumor growth was assessed as described previously (1,9). When tumor diameter reached approximately 0.5 cm, the animals were randomized into four groups of treatment, with at least five animals in each group: control (vehicle-treated), lapatinib alone, CQ and the combination of lapatinib + CQ. Lapatinib was resuspended in an aqueous solution containing 0.1% Tween 80 (Sigma-Aldrich) and 0.5% Hypromellose (Sigma-Aldrich), and delivered orally at 100 mg/kg once daily. CQ was solubilized in sterile 1X phosphate-buffered saline (PBS) and administered intraperitoneally at 50 mg/kg once daily. Mice were weighed and tumor growth was monitored twice a week. Tumor size was measured using a caliper and the tumor volume was calculated using the formula $L \times W^2 \times (\pi/6)$, where L is the length and W, the width of the tumor xenografts. At the end of the experiment, the mice were sacrificed, and the tumors extracted and processed for histological evaluation. A hematoxylin and eosin (H&E) staining and GSDMB and PCNA immunohistochemical analysis was performed in all of the tumors as mentioned above.

**Immunohistochemistry analysis.** Human and mouse tumor samples were collected, fixed in 4% formaldehyde and embedded in paraffin. Three µm thick sections were stained with H&E solutions. Primary antibodies used are listed in **Supplementary Table S5**. Briefly, tumor sections were deparaffinized using the BOND Dewax Solution (Leica Biosystems) and rehydrated with serial passage through changes of decreasing ethanol solutions. Immunostaining was performed on a BOND $RX^m$ autostainer (Leica Biosystems) using BOND Epitope Retrieval Solutions (Leica Biosystems) and the BOND Polymer Refine Detection kit (Leica Biosystems), following the manufacturer's instructions. Stained slides were counterstained with hematoxylin and coverslipped for obtaining the images with an Aperio CS2 scanner (Leica Biosystems) that were analyzed with the Aperio eSlide Manager software (Leica Biosystems). Immunohistochemical analyses in tumor sections of GSDMB and LC3B were performed according to stablished protocols(2,4). Primary antibodies used are listed in **Supplementary Table S5**. A semiquantitative scoring of each marker based on staining intensity and the percentage of stained tumor cells was applied by two independent observers, blinded to clinical data.

**Prediction tools.** iLIR database (https://ilir.warwick.ac.uk/) was used to detect LIR motifs on the full-length GSDMB protein (10). With respect to the *in silico* protein interaction prediction, the GSDMB and LC3B protein structures were predicted using the swiss-model algorithm (11). 3D structures of LC3B and GSDMB were obtained from Uniprot data bases (accession number Q9GZQ8 and Q8TAX9, respectively). Then, the protein-protein complex was deduced using the algorithm HawkDock (12). This method allows selecting the possible protein complex in function of predicted Gibbs energy. The structure with the least Gibbs energy, in this case, -2 Kcal/mol, was selected. The predicting interacting residues of the protein complex were analyzed using the InterProSurf (13) and PPcheck (14) webservers.

**SUPPLEMENTARY TABLES**

**Supplementary Table S1.** Summary clinic-pathological data of the HER2 gastric carcinoma cohort*.

| | †N (%) |
|---|---|
| **GSDMB OVER-EXPRESSION** | |
| LOW | 13/31 (41,9) |
| HIGH | 18/31 (58.1) |
| **LC3B PUNCTA EXPRESSION** | |
| LOW | 13/26 (50.0) |
| HIGH | 13/26 (50.0) |
| **RELAPSE#** | |
| NO | 13/32 (40.6) |
| YES | 19/32 (59.4) |
| **DISTANT METASTASIS** | |
| NO | 12/32 (37.5) |
| YES | 20/32 (62.5) |

The data refer to the available cases for each parameter. †n (%), number of analyzed cases and (percentage). #**Relapse,** local or distant recurrence.

**Supplementary Table S2.** *In vivo* acute toxicity results in zebrafish.

| Anticancer drug | Concentration, μM | | | |
|---|---|---|---|---|
| | **LC10** | **LC50** | **NOEC*** | **LOEC*** |
| **Lapatinib** | 31.2 | 35.1 | <30.0 | <30.0 |
| **Chloroquine** | 74.1 | 116.4 | 100 | 50 |

*LOEC lowest observed effect concentration, NOEC no-observable effect concentration values

**Supplementary Table S3.** Summary clinic-pathological data of the adjuvant HER2+ breast carcinoma cohort*.

| | †N (%) |
|---|---|
| **GSDMB OVER-EXPRESSION** | |
| LOW | 19/32 (59.4) |
| HIGH | 13/32 (40.6) |
| **LC3B PUNCTA EXPRESSION** | |
| LOW | 14/32 (43.7) |
| HIGH | 18/32 (56.3) |
| **RELAPSE#** | |
| NO | 13/32 (40.6) |
| YES | 19/32 (59.4) |

*HER2 breast cancer cohort previously reported (Hergueta-Redondo, 2016). The data make reference to the available cases for each marker. †n (%), number of analyzed cases and (percentage). #**Relapse,** local or distant recurrence.

**Supplementary Table S4**. List of primers used for qRT-PCR and cloning.

Myc-epitope sequence: bold labelled

| Gene | Forward | Reverse |
|---|---|---|
| **GSDMA** | 5’-AAACCTTCCCTCCTGGAGAA-3’ | 5’-CCCTTGTCTAGAGCCCCTTC-3’ |
| **GSDMB** | 5’-TGGCTACCCTTGAAAACAGG-3’ | 5’-ATATTGCCGGTCGCTTTTC-3’ |
| **GSDMC** | 5’-AGCTCAGGTCATTTGGATGG-3’ | 5’-GCACCATTATGGCTTCAAGG-3’ |
| **GSDMD** | 5’-GCGTCAGGTTGCAGTTTCAC-3’ | 5’-CGGACTACCCGCTCAAAGG-3’ |
| **GSDME** | 5’-GAAGATCACGACGATGCAGA-3’ | 5’-GAAGGCAGAACTCGAACTGG-3’ |
| **HER2** | 5’-CGAGTGTCAGCCTCAAAACA-3’ | 5’-TCATCCAGATCCACACAGGA-3’ |
| **GAPDH** | 5’-TGCCATCAATGACCCCTTC-3’ | 5’-ATTTGCCATGGGTGGAATC-3’ |
| **GB$^{92\text{-}416}$-myc** | 5’-GCGAATTCATGTCTACGGGCG AGCTGAT-3’ | 5’-GCCCCACTAGCGTGAGCAGCGAACA AAAACTCATCTCAGAAGAGGATCTGTAGT CTAGA-3’ |

**Supplementary Table S5.** List of primary antibodies used for western blot (WB), immunofluorescence (IF), immunoprecipitation (IP), and immunohistochemistry (IHC).

| Antibody | Species* | Source | Identifier | WB/IF/IP/IHC |
|---|---|---|---|---|
| **anti-ATG5 (C-terminal)** | rPab | Sigma-Aldrich | A0731 | 1:500/-/-/- |
| **anti-Caspase-3 (Clone 31A1067)** | mMab | Santa Cruz Biotechnology | sc-56053 | 1:250/-/-/- |
| **anti-GSDMB** | mMab | Hergueta-Redondo et al., 2016 | --- | 1:250/-/-/1:200 |
| **anti-GAPDH (Clone 6C5)** | mMab | Sigma-Aldrich | CB1001 | 1:50000/-/-/- |
| **anti-LC3B** | rPab | Sigma-Aldrich | L7543 | 1:1000/1:10/1:50/1:2000 |
| **anti-MAP LC3beta** | mMab | Santa Cruz Biotechnology | sc-376404 | -/-/-/1:200 |
| **anti-Myc-Tag (Clone 9B11)** | mMab | Cell Signaling Technology | 2276 | 1:1000/1:100/1:200/- |
| **anti-Myc-Tag (Clone 71D10)** | mPab | Cell Signaling Technology | 2278 | 1:1000/-/-/- |
| **anti-HER2 (Clone 44E7)** | mMab | Cell Signaling Technology | 2248 | 1:1000/-/-/- |
| **anti-Phospo-HER2 (Tyr1221/1222)** | rMab | Cell Signaling Technology | 2243 | 1:1000/-/-/- |
| **anti-SQSTM1/p62 (Clone D5E2)** | rMab | Cell Signaling Technology | 8025 | 1:1000/-/-/- |
| **PCNA (clone p10)** | mMab | EMD MILLIPORE | MAB424R | -/-/-/1:10000 |
| **HRP-conjugated secondary antibody** | mMab | GE Healthcare Life Sciences | NA931 | 1:5000/-/-/- |
| **HRP-conjugated secondary antibody** | rPab | GE Healthcare Life Sciences | NA934 | 1:5000/-/-/- |
| **Alexa Fluor 488-labeled** | mMab | Invitrogen | A-11029 | -/1:1000/-/- |

| | | | | |
|---|---|---|---|---|
| **secondary antibody** | | | | |
| **Alexa Fluor 546-labeled secondary antibody** | rPab | Invitrogen | A-11035 | -/1:1000/-/- |
| **Alexa Fluor 488-labeled secondary antibody** | rPab | Invitrogen | A-11034 | -/1:1000/-/- |

* rPab: rabbit polyclonal antibody, mMab: Mouse monoclonal antibody, rMab: rabbit monoclonal antibody

**SUPPLEMENTARY FIGURES S1-7.**

A

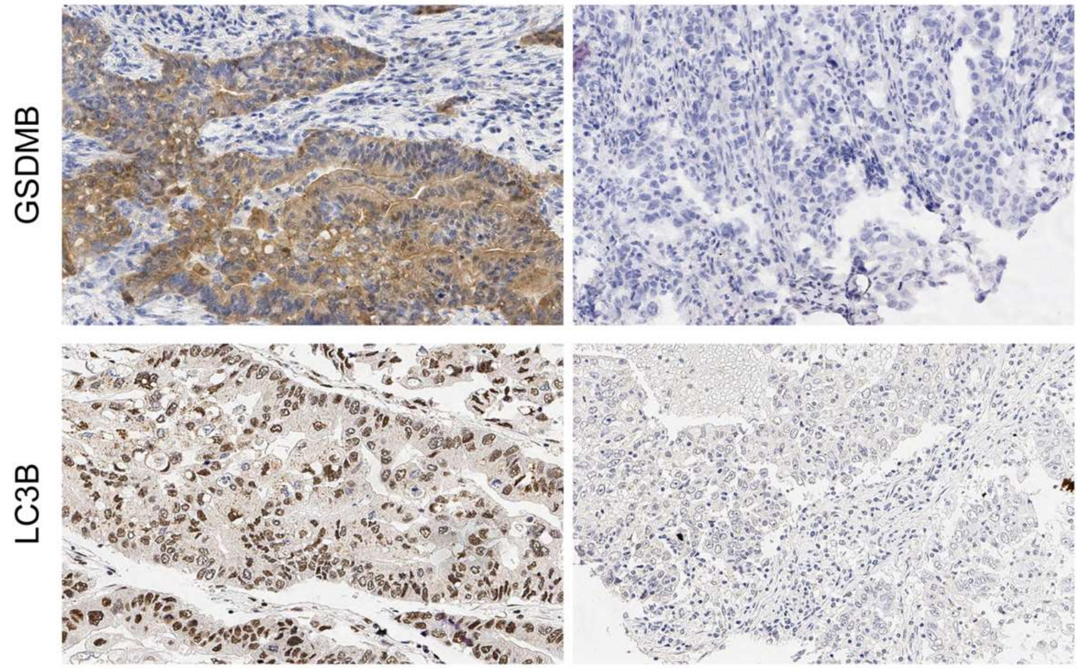


B

| | GSDMB | |
|---|---|---|
| | LOW n (%) | HIGH n (%) |
| **Metastasis** | | |
| Negative | 8/13 (61.5) | 4/18 (22.2) |
| Positive | 5/13 (38.5) | 14/18 (77.8) |
| | | *p=0.032* |
| **Relapse** | | |
| Negative | 8/13 (61.5) | 5/18 (27.8) |
| Positive | 5/13 (38.5) | 13/18 (72.2) |
| | | *p=0.060* |
| **LC3B puncta expression** | | |
| Low | 7/9 (77.8) | 5/15 (33.3) |
| High | 2/9 (22.2) | 10/15 (66.7) |
| | | *p=0.045* |

| | Relapse | |
|---|---|---|
| | NEGATIVE n (%) | POSITIVE n (%) |
| **GSDMB & LC3B coexpression** | | |
| No | 8/14 (57.1) | 1/10 (10.0) |
| Yes | 6/14 (42.9) | 9/10 (90.0) |
| | | *p=0.024* |

**Supplementary Figure S1.** GSDMB over-expression is associated with metastasis, relapse and LC3B expression in HER2 gastric tumors. **A,** Representative immunohistochemistry images of GSDMB and LC3B positive (left) and negative expression (right) in a cohort of HER2 gastric carcinomas. **B,** Correlation between GSDMB and metastasis, relapse and LC3B expression as well as GSDMB/LC3B co-expression with relapse in HER2 gastric carcinomas. Differences were compared by Fisher's exact test.

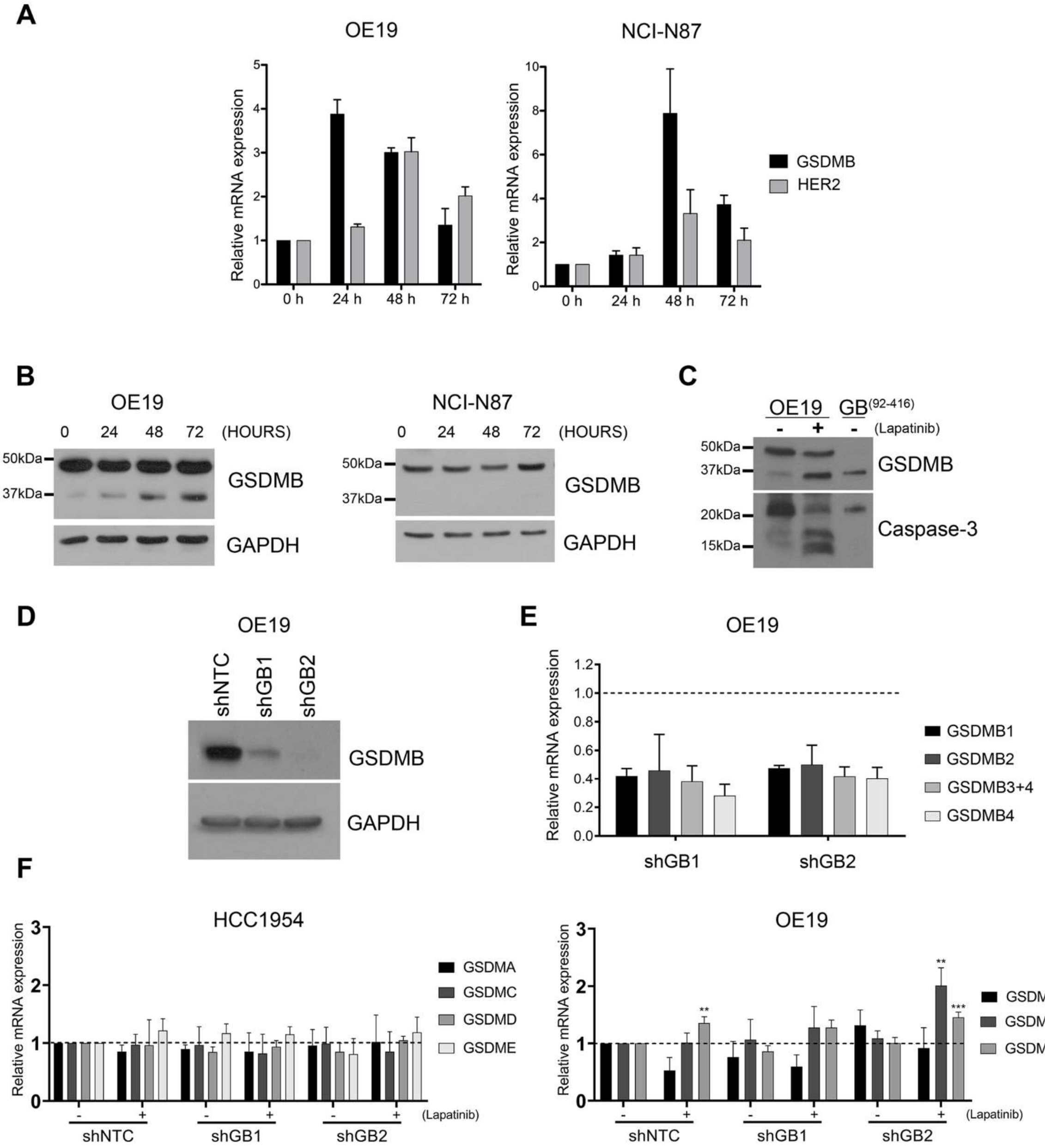


**Supplementary Figure S2.** Upregulation of GSDMB in response to trastuzumab and expression of different GSDMs in GSDMB-silenced cells after treatment with lapatinib. **A** and **B,** Relative mRNA (**A**) and protein levels (**B**) of GSDMB and HER2 in OE19 and NCI-N87 cells treated with IC50 of trastuzumab (400 µg/ml in both cases) at different time points. **C,** GSDMB and caspase-3 analyses on OE19 control, treated with lapatinib and over-expressing $GB^{92-416}$ cells. $GB^{92-416}$ is the C-terminal fragment previously described (29) and generated after the caspase 3 processing. **D** and **E,** GSDMB protein (the four translated GSDMB isoforms, **D**) and mRNA (**E**) levels were stably decreased in OE19 cells by two specific shRNAs (shGB1 and shGB2), in comparison with the control (shNTC; dotted line). **F,** GSDMB-silencing effect in the mRNA levels of the other Gasdermin genes (GSDMA, GSDMC, GSDMD and GSDME) in HCC1954 and OE19 cells treated with or without lapatinib (2 µM and 0.7 µM, respectively, for 72 h). Statistical significance was determined by two-tailed unpaired *t*-test (**P* < 0.05; ***P* < 0.01; ****P* < 0.001). Gene expression was normalized to GAPDH levels. Data are shown as the mean ± s.e.m. Three independent experiments with similar results were performed.

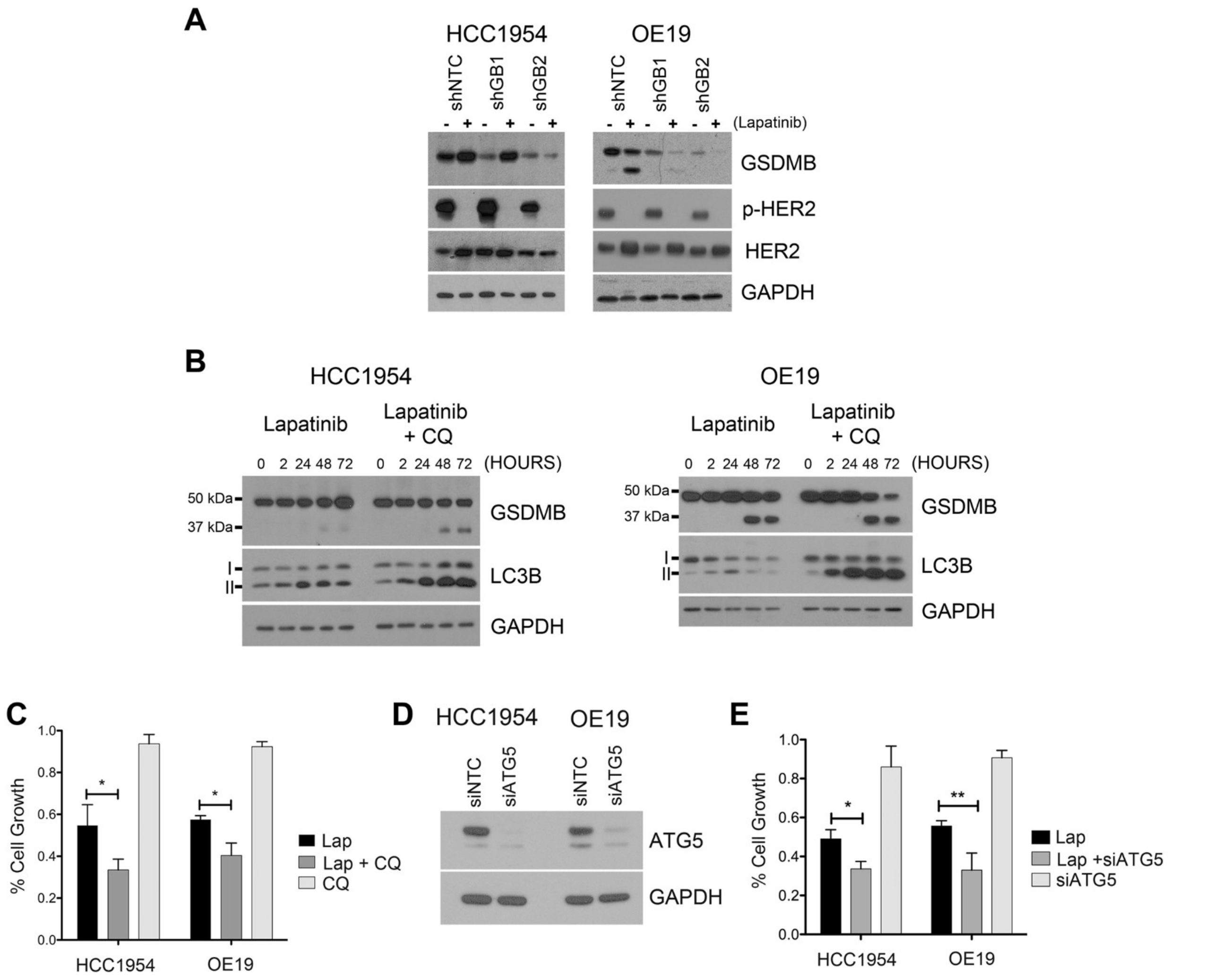


**Supplementary Figure S3.** GSDMB induces pro-survival autophagy in response to lapatinib. **A,** GSDMB-silencing effect on the levels of HER2 receptor in HCC1954 and OE19 cells treated with or without IC50 of lapatinib (2 µM and 0.7 µM, respectively, for 72 h). **B,** GSDMB and LC3B protein levels in HCC1954 and OE19 cells treated with lapatinib (2 µM and 0.7 µM, respectively) or lapatinib + CQ (10 µM and 50 µM, respectively) at different time points. **C,** The cytotoxic effect of the treatment with lapatinib and/or CQ in HCC1954 and OE19 cells was evaluated by cell viability assays. **D,** ATG5 expression was effectively decreased in HCC1954 and OE19 cells by a specific siRNA (siATG5), compared to the control (siNTC). **E,** Similarly, to CQ treatment, the autophagy inhibition by siATG5 improves the effect of lapatinib in HCC1954 and OE19 cells, measured by cell viability assays. Three independent experiments with similar results were carried out. In (**C** and **E**), statistical significance was determined by two-tailed unpaired *t*-test (*$P$ < 0.05; **$P$ < 0.01). Data are shown as the mean ± s.e.m. CQ, chloroquine. NTC, non-targeting control.

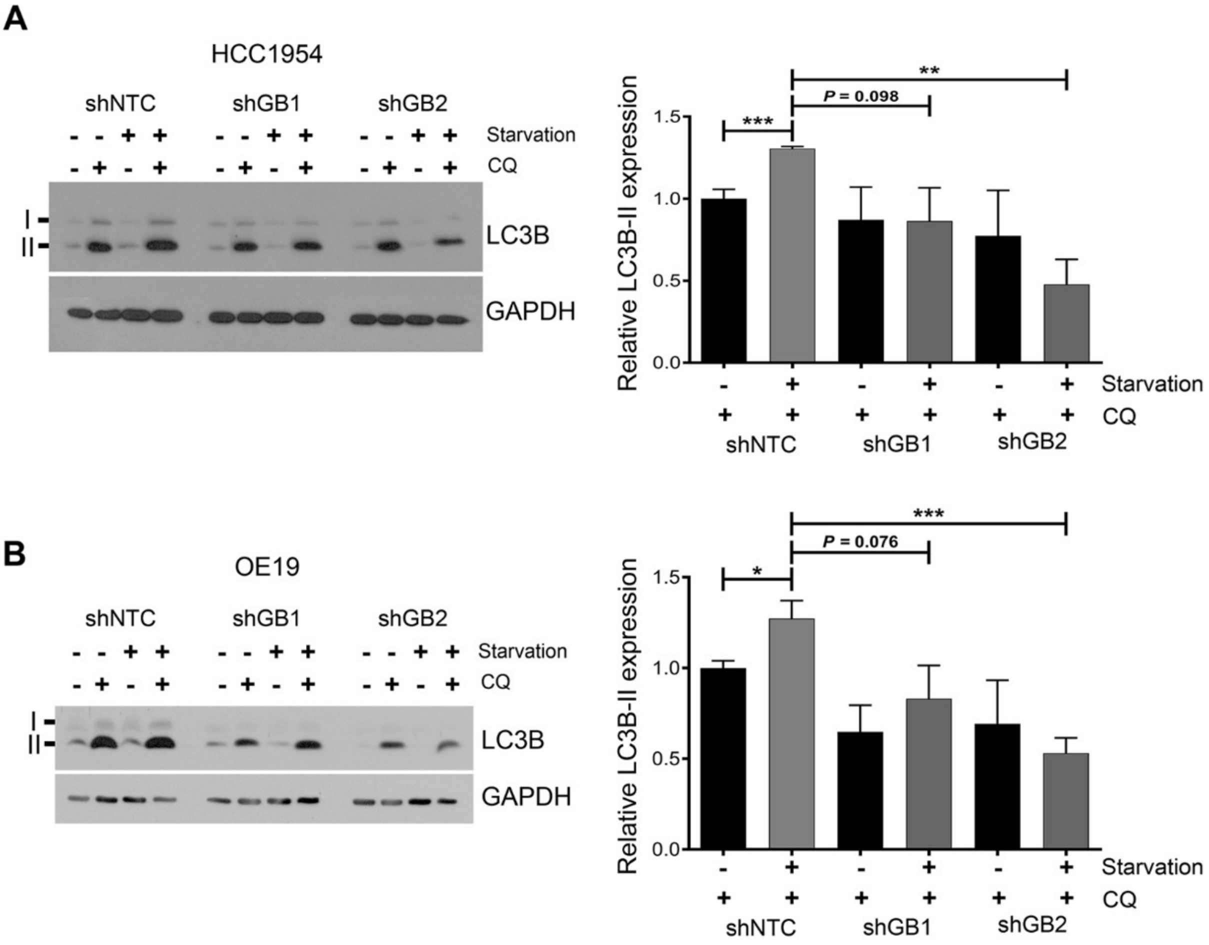


**Supplementary Figure S4.** GSDMB-high cells show an increased autophagic flux in response to starvation. **A** and **B,** GSDMB and LC3B protein levels in shNTC, shGB1 and shGB2 HCC1954 (**A**) and OE19 (**B**) cells treated with IC50 of lapatinib (2 μM and 0.7 μM, respectively) and/or serum starved (0.01% fetal bovine serum) for 72 h. Quantification of LC3B-II expression was carried out by densitometric scanning and normalized to GAPDH expression. Statistical significance was determined by two-tailed unpaired t-test (**P* < 0.05; ***P* < 0.01; ****P* < 0.001; ns, nonsignificant). Data are shown as the mean ± s.e.m. Three independent experiments with similar results were carried out. NTC, non-targeting control.

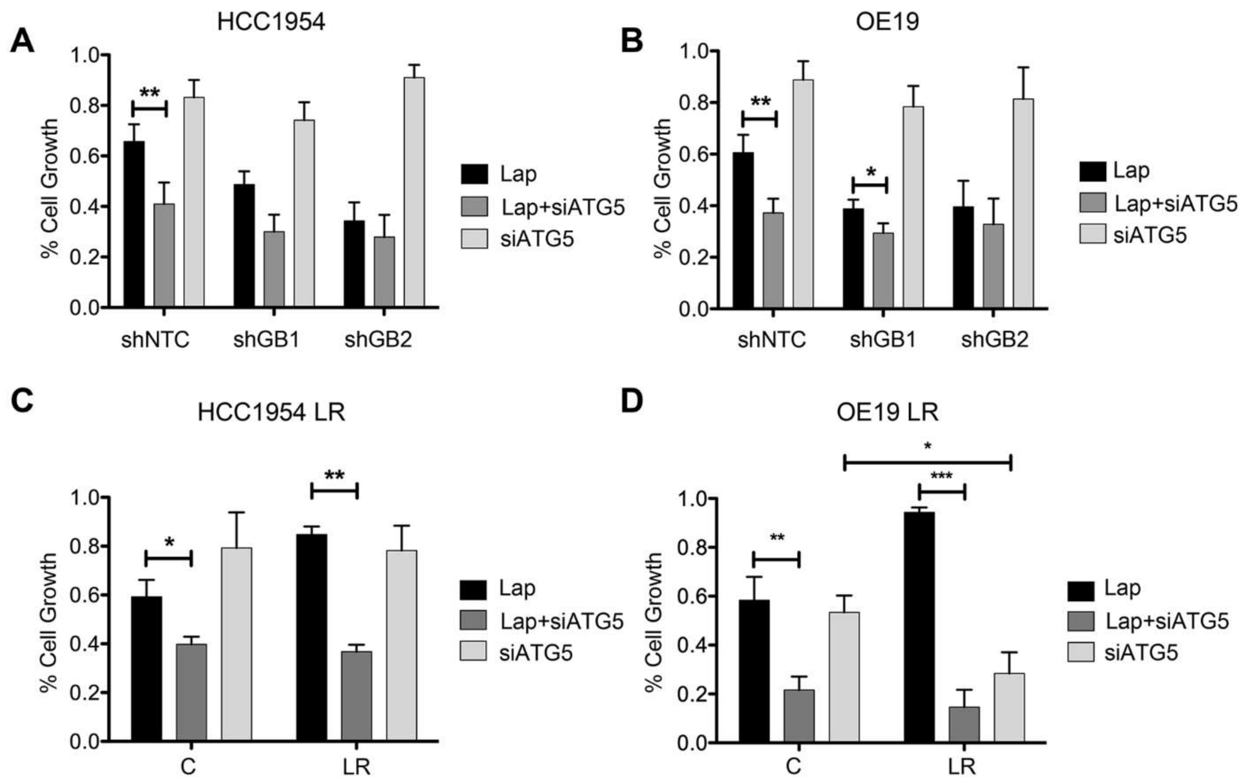


**Supplementary Figure S5.** ATG5-silencing renders GSDMB-high cells more sensitive to lapatinib. **A-D,** The cytotoxic effect of the treatment with lapatinib plus/or ATG5-silencing was evaluated by cell viability assays in shNTC, shGB1 and shGB2 HCC1954 (**A**) and OE19 cells (**B**), as well as HCC1954 LR (**C**) and OE19 LR cells (**D**). Statistical significance was determined by two-tailed unpaired t-test (*P < 0.05; **P < 0.01; ***P < 0.001). Data shows the mean ± s.e.m of three independent experiments with similar results. NTC, non-targeting control.

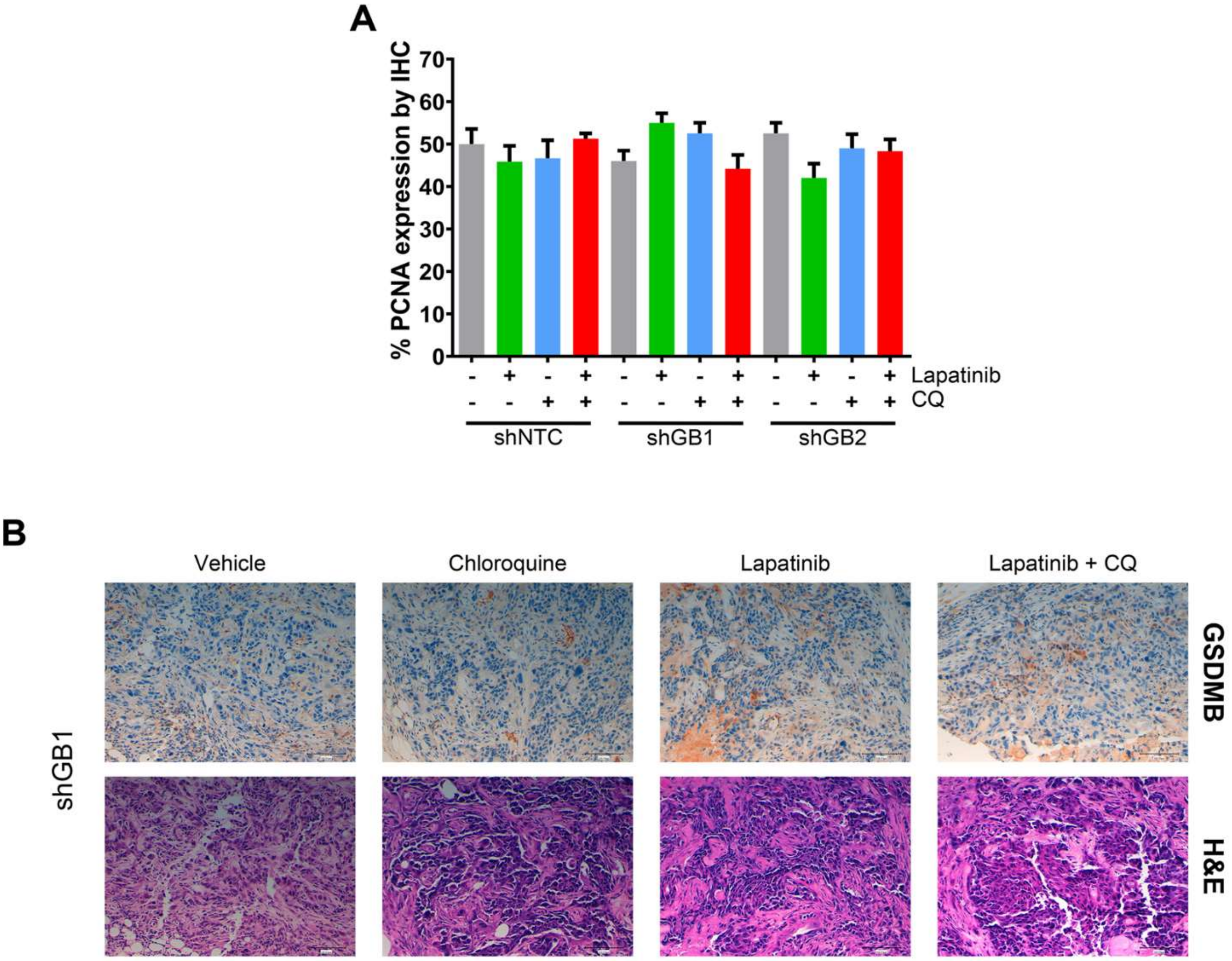


**Supplementary Figure S6.** Immunohistochemical and histological analysis in orthotopic tumor xenografts. **A,** Quantification of PCNA immunohistochemical expression on HCC1954-mCherry-luc control (shNTC), or GSDMB-silenced cells (shGB1 and shGB2) xenografted tumors treated with lapatinib, Chloroquine (CQ), or a combination of both. **B,** Representative images of GSDMB immunohistochemical analysis and hematoxylin and eosin staining in shGB1 xenografted tumors, treated with the different therapeutic strategies indicated in (**A**) (scale bar, 100 μm). NTC, non-targeting control.

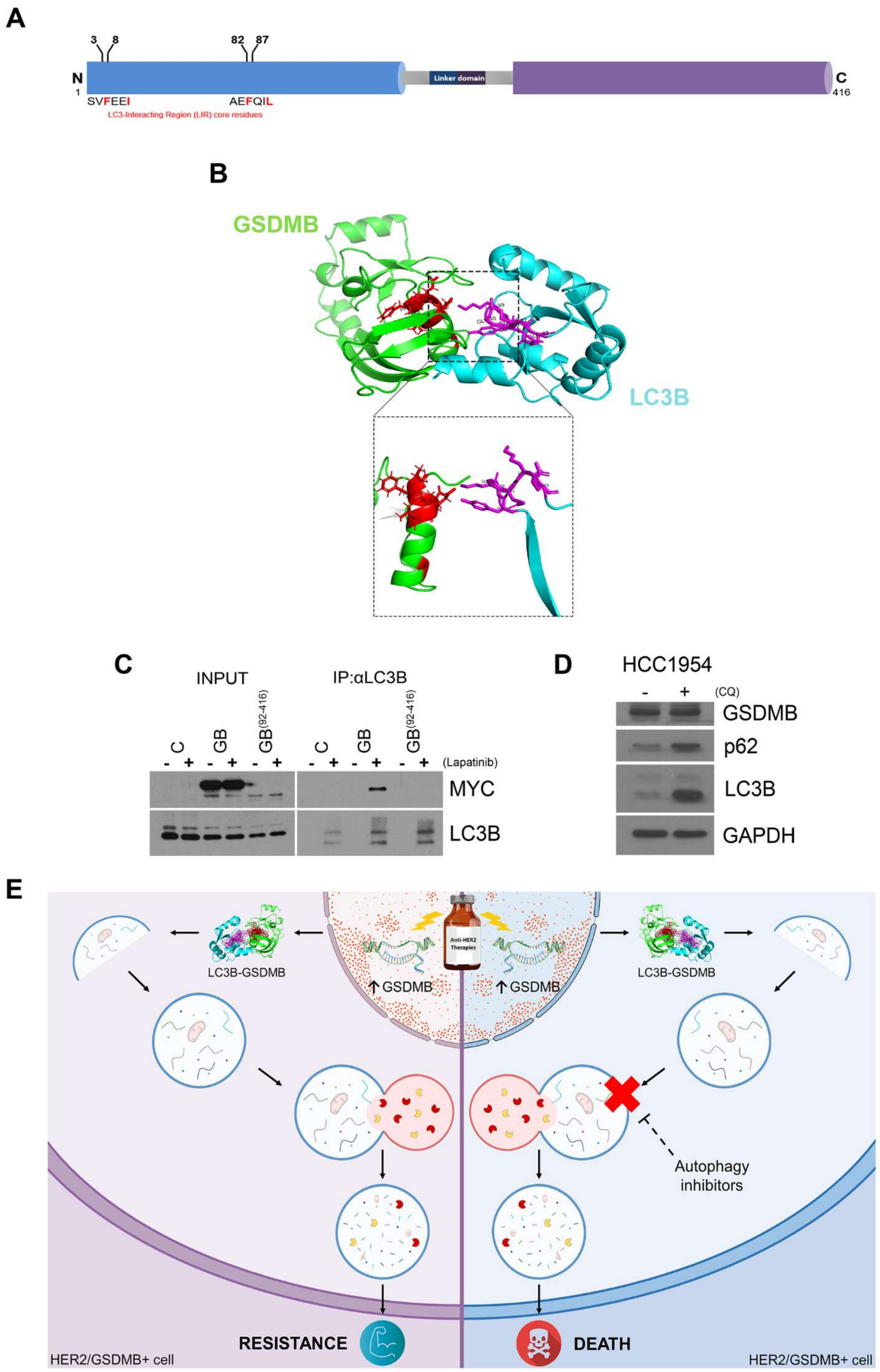


**Supplementary Figure S7.** Potential interaction sites between GSDMB and LC3B through LIR domains. GSDMB is not degraded during autophagy. **A,** Schematic representation of GSDMB showing the LIR motifs (core residues in red) located on the N-terminal region. **B,** *In silico* protein interaction prediction using HawkDock algorithm (39) identifies a potential interaction between the N-terminal region of LC3B (blue) and the LIR1 motif within the GSDMB N-terminal domain (green). Inset magnifies the predicting interacting region (GSDMB, red and LC3B, pink). 3D structures were obtained from Uniprot databases (Q9GZQ8: L3CB and Q8TAX9: GSDMB). **C,** Co-immunoprecipitation assay of LC3B-GSDMB interaction after autophagy induction with lapatinib in HCC1954 cells exogenously overexpressing full length GSDMB (GB), a construct lacking the LIR motifs ($GB^{(92-416)}$) or control (C; empty vector) cells, using an anti-LC3B antibody. **D,** GSDMB, p62 and LC3B protein levels in HCC1954 cells treated for 72 h with or without chloroquine (10 μM). **E,** Representative scheme of the proposed mechanism of action of GSDMB as mediator of protective autophagy response. It is highlighted the dualistic model on HER2/GSDMB over-expressing tumors behavior after the treatment anti-HER2 therapy (resistance phenotype) or its combination with autophagy inhibitors (sensible phenotype).
